# Alternative splicing induced by bacterial pore-forming toxins sharpens CIRBP-mediated cell response to *Listeria* infection

**DOI:** 10.1101/2023.01.18.524578

**Authors:** Morgane Corre, Volker Boehm, Vinko Besic, Anna Kurowska, Anouk Viry, Ammara Mohammad, Catherine Sénamaud-Beaufort, Morgane Thomas-Chollier, Alice Lebreton

**Affiliations:** Group Bacterial infection, response & dynamics, Institut de biologie de l’ENS (IBENS), École normale supérieure, CNRS, Inserm, Université PSL, 75005 Paris, France; Institute for Genetics, University of Cologne, Cologne, Germany; Center for Molecular Medicine Cologne (CMMC), University of Cologne, Cologne, Germany; GenomiqueENS, Institut de Biologie de l’ENS (IBENS), École normale supérieure, CNRS, INSERM, Université PSL, 75005 Paris, France; INRAE, Micalis Institute, 78350 Jouy-en-Josas, France

## Abstract

Cell autonomous responses to intracellular bacteria largely depend on gene expression reorganization. To gain isoform-level resolution into these regulations, we combined long- and short-read transcriptomic analyses of the response of intestinal epithelial cells to infection by the foodborne pathogen *Listeria monocytogenes*. Among the most striking isoform-based regulations, expression of the cellular stress response regulator CIRBP (cold-inducible RNA-binding protein) and of several SRSFs (serine/arginine-rich splicing factors) switched from canonical transcripts to nonsense-mediated decay-sensitive isoforms by inclusion of “poison exons”. We showed that damage to host cell membranes caused by bacterial pore-forming toxins (listeriolysin O, perfringolysin, streptolysin, or aerolysin) led to the dephosphorylation of SRSF proteins *via* the inhibition of the kinase activity of CLK1, thereby driving CIRBP alternative splicing. CIRBP isoform usage was found to have consequences on infection, since selective repression of canonical CIRBP reduced intracellular bacterial load while that of the poison exon-containing isoform exacerbated it. Consistently, CIRBP-bound mRNAs were shifted towards stress-relevant transcripts in infected cells, with increased mRNA levels or reduced translation efficiency for some targets. Our results thus generalize the alternative splicing of CIRBP and SRSFs as a common response to biotic or abiotic stresses by extending its relevance to the context of bacterial infection.

## Introduction

In a fluctuating environment or upon perturbation of tissue homeostasis, cells can undergo a broad variety of damaging conditions that may impair their integrity and function. In response to the action of damaging agents, stress-sensitive pathways coordinate multiple cellular mechanisms of response that include the control of cell cycle checkpoints, the reallocation of resource between major anabolic and catabolic pathways, the protection against macromolecular damage and scavenging of damaged components, the launching of pro-inflammatory responses, or eventually cell death (1, 2). Among key regulators involved in orchestrating cellular stress responses, CIRBP has been previously described as a central stress-response protein. Although it was initially identified for its role in cold-stress adaptation, it also responds to a variety of other non-physiological challenges such as hypoxia, DNA damage response, glucose deprivation, osmotic-, oxidative- or heat-stresses (3–5).

Downstream of stress-sensing, the expression of CIRBP has been shown to be regulated not only in abundance, but also via the production of alternative transcripts (6). Accordingly, 46 splice variants of human CIRBP are annotated in the Ensembl database version 109.38 (<u>ENSG00000099622</u>). The mechanisms of CIRBP isoform conversion have been previously investigated in the context of mild cold- and heat shocks, revealing a fine tuning of CIRBP pre-mRNA splicing into alternative isoforms (7–9). Even a moderate increase in temperature results in the inhibition of the activity of the kinase CLK1, leading to the dephosphorylation of serine/arginine-rich splicing factors (SRSFs, also known as SR proteins). In turn, the loss of activity of SR proteins drives the splicing of CIRBP pre-mRNA in favor of unstable isoforms that are sensitive to the nonsense-mediated decay (NMD) pathway of RNA degradation.

The CIRBP protein itself is activated and migrates to cytoplasmic stress granules, where it regulates a number of cellular pathways including cell cycle and inflammation, via its RNA binding activity and ability to modulate the stability and translation of its targets (3, 10, 11). The modulatory role of CIRBP on inflammation is complex and context-dependent; while it acts as a mediator of tumor-promoting inflammation, it dampens inflammation in the resolution phase of wound healing (10). It was also reported that in Huh7 cells from liver hepatoma, viral infections resulted in the loss of *N*^6^-methyladenosine (m^6^A) in CIRBP 6^th^ exon, located in the 3’ region of CIRBP coding sequence (CDS) (12). This change in CIRBP pre-mRNA methylation led to alternative splicing (AS) of the transcript and as a consequence, to the loss of a long, cytoplasmic protein-coding isoform. Expression of both this long form of CIRBP and the main, shorter isoform, which displayed a predominantly nuclear distribution, had a proviral effect. In addition to intracellular roles of CIRBP as an RNA-binding protein, during haemorrhagic shock or sepsis CIRBP can be released in the circulation, where it acts as a damage-associated molecular pattern (DAMP) that promotes inflammatory responses (13). At least part of the inflammatory effects and tissue injury caused by extracellular CIRBP (eCIRBP) during sepsis relies on its ability to induce neutrophil extracellular traps (14).

While CIRBP functions have been investigated in various stress conditions including viral infections and sepsis, its regulation and cellular roles have not been documented in the context of bacterial infections. Yet, bacterial challenges have been reported to provoke several types of stresses that are sensed by host cells (15–17), including during infection by the foodborne pathogen *Listeria monocytogenes* (*Lm).* This facultative intracellular bacterium is responsible for listeriosis in cattle and humans, a rare but life-threatening disease in immunocompromised individuals and in the elderly, which can cause death from meningitis, encephalitis or septic shock (18). Maternofetal transmission of listeriosis can also cause miscarriage, stillbirth, or debilitating neonatal meningitis. A number of studies have reported that *Lm* infection is accompanied by cellular damages that cause potent cellular stress responses such as the activation of the integrated stress response pathway (19–21). The major cause of *Lm*-triggered stress to its host cell is the permeation of cellular membranes by a cholesterol-dependent cytolysin, listeriolysin O (LLO) (22–25).

Here, using a combined approach based on short- and long-read sequencing of the transcriptome of intestinal epithelial cells upon infection by *Lm*, we show that AS events alter the expression of a number of cellular genes involved in host RNA processing and post-transcriptional regulation, including splicing factors and CIRBP. We identify that the pore-forming activity of LLO is responsible for CIRBP isoform conversion towards NMD-sensitive transcripts, *via* the inhibition of the activity of CLK1 kinase and of SR proteins. We observe an opposite effect of the two classes of CIRBP isoforms, and identify variations in the sets of CIRBP targets in infected *versus* uninfected cells as well as in their expression. Altogether, our findings generalize the mechanism of CIRBP-based stress responses to the context of a bacterial infection.

### Material and methods

#### Biological resources

LoVo cells, a human intestinal epithelial cell line originating from a colon adenocarcinoma (male, ATCC CCL-229) were used as a main host cell model. The Hep G2 line of hepatocellular carcinoma (ATCC HB-8065) and THP-1 line of human monocytes (ATCC TIB-202) were used to generalize the effects of LLO on CIRBP AS. Cells were maintained in their respective formulated media, supplemented with 10% heat-inactivated fetal bovine serum (PANBiotech, cat#p30-3306) in a humidified incubator at 37°C and 5% CO_2_. Formulated media were: for LoVo, DMEM, low glucose, GlutaMAX supplement, pyruvate medium (ThermoFisher Scientific, cat#21885025); for Hep G2, Eagles’ MEM (Corning, cat#10009CV); for THP-1, RPMI 1640 medium (ThermoFisher Scientific, cat#21875034). THP-1 cells were activated with 100 μg/mL of phorbol 12-myristate-13-acetate (PMA) to induce their differentiation into macrophages, 48 h before each experiment.

The bacterial source strains used for this work (Table S1) were *Listeria monocytogenes* LL195 as a main infection model and *Escherichia coli* NEB5α (New England BioLabs) for protein purification. *Listeria innocua* [*inlA*] and *L. monocytogenes* LL195 Δ*hlyA* were used for comparative purposes in infection assays. *E. coli* Top10(DE3) [pET29b-LLO-6His], *E. coli* Top10 [pET29b-LLO_C484A_-6His], *E. coli* Top10 [pET29b-LLO_W492A_-6His], *E. coli* Top10 [pET29b-LLO_Y206A_-6His], *E. coli* DH5α [pTrcHisA-PFO], *E. coli* DH5α [pTrcHisA-SLO] and *E. coli* NEB5α [pET22b-PA-6His] were used as sources for the purification of pore-forming toxins. All strains were grown at 37°C under shaking at 180 rpm in Luria Bertani (LB) medium for *E. coli*, or in brain heart infusion (BHI) for *Lm* and *L. innocua* strains. Whenever required, media were supplemented with antibiotics for plasmid selection (chloramphenicol, 7 μg/μL for *Lm* harboring a pAD vector and *inlA*-expressing *L. innocua*; kanamycin, 30 μg/μL for *E. coli* Top10 harboring pET29b vectors; and ampicillin, 100 μg/μL for *E. coli* DH5α and *E. coli* NEB5α harboring pTrcHisA and pET22b vectors).

#### Culture, transfection and infection of epithelial cells

Infections were performed in LoVo cells. Cells were between passage 7 and passage 15 before seeding and were grown to 80-85% confluence for 48 h prior to infection. When needed, cells were transfected 48 h before infection with 6 pg siRNAs against CIRBP-ORF, CIRBP-201 and CIRBP-210, UPF1 or scrambled siRNAs (Table S2A) using RNAiMAX in 12-well format as per manufacturer’s recommendations (reverse transfection procedure). The decreased abundance of the corresponding proteins was confirmed by RT-qPCR and western blot.

One colony of bacteria was grown for about 16 h until they reached stationary phase (optical density at 600 nm — OD_600_ of 2 to 3) in 5 mL of BHI medium at 37°C. Bacteria were washed twice with PBS and added to the cell monolayer in culture flasks (for RNA-seq or RNA Immunoprecipitation (RIP)) or in 12-well plate format at the appropriate MOI (see Table S1A for details), by diluting bacteria to the appropriate density in serum-free DMEM. The cell culture flasks or plates were centrifuged at 200 × g for 1 min and then incubated at 37°C and 5% CO_2_ for 1 h. Cells were then washed with PBS containing 25 μg/mL gentamicin, after which fresh medium containing 25 μg/mL gentamicin and 10% FBS was added. Infection was allowed to proceed until specific timepoints (2, 5 or 10 hpi, or uninfected for time 0), then culture plates were snap-frozen in liquid nitrogen and stored at −80°C until further use (for RNA or protein extraction or for RIP). Alternatively, for infection quantification, cells were washed in PBS and trypsinized for counting using a LUNA II automated cell counter, then centrifuged at 700 x g for 5 min. Cell pellets were resuspended in 500 μL ice-cold water and lysed by passing several times through a 26G needle. Cells lysates were diluted in water and plated on BHI before overnight incubation at 37°C. Colony-forming units (CFU) were counted and normalized to cell counts.

#### Immunoblot assays

Total cell lysate was prepared by adding RIPA (20 mM Tris-HCl pH 7.5, 150 mM NaCl, 1 mM EDTA, 1 mM EGTA, 1% IGEPAL, 1% Sodium Deoxycholate, 50 μg/μL PMSF) supplemented with protease inhibitor cocktail (Bimake, cat#B14011) and when needed, phosphatase inhibitor cocktail (Sigma, cat#P5726) directly to the cell monolayer. After scraping of the monolayer, the lysate was transferred to an Eppendorf tube, incubated on ice for 10 min with intermittent vortex mixing, and then centrifuged for 10 min at 4°C at maximum speed. The supernatant was transferred to a fresh Eppendorf tube and the protein concentration was assessed using Bradford reagent (Abcam, cat#ab119216) and stored at -80°C when not used immediately. 40 μg of proteins were diluted in Laemmli sample buffer (SB 1X), after which samples were heated at 95°C for 5 min and either stored at −80°C or used directly. Samples were separated by SDS-PAGE on a 12%-acrylamide gel and transferred to Amersham Protran 0.2 μM nitrocellulose membranes using a Mini Trans-Blot cell (Bio-Rad) for 1 h at 100 V at 4°C in chilled transfer buffer (Tris-Base 3 g/L, glycine 1.4 g/L, 20% ethanol). Membranes were blocked for 30 min in PBS or TBS containing 0.1% of Tween 20 (PBS-T and TBS-T, respectively) and 5% milk or bovine serum albumin (BSA), according to antibody manufacturer’s recommendations. Primary antibodies were added to the blocking solutions at the recommended dilution before overnight incubation at 4°C. Anti-CIRBP (Proteintech, cat#10209-2-AP, 1:500 dilution); anti-β-Tubulin (Biolegend, cat#903401, 1:2, 000 dilution); anti-GAPDH (Cell Signaling Technologies, cat#2118, 1/1, 000 dilution); anti-SR proteins (Millipore, cat#MABE126, 1:2, 000 dilution); anti-phosphoepitope SR proteins clone1H4 (Millipore, cat#MABE50, 1:2, 000 dilution) were used as primary antibodies. Membranes were washed three times in PBS-T or TBS-T and incubated with the corresponding secondary antibody (Bethyl anti mouse or rabbit IgGs coupled to HRP, cat#A120101P and A90-116P, respectively) at a 1:50, 000 dilution in the same buffer for 2 h at room temperature. Signals were revealed using Pierce ECL Plus Western Blotting Substrates (ThermoFisher Scientific, cat#32106 and #32132, respectively) on an Las4000 imager (Amersham).

#### RNA extraction and RT-qPCR

From a 12-wells plate format, total RNA was extracted by adding 300 μL of TriReagent (MRC, cat#TR118) directly onto the cell monolayer and following the extraction procedure according to the manufacturer’s instructions. After ethanol precipitation, RNA was resuspended in 44 μL of RNase-free water, to which 5 μL of Turbo DNase 10X Buffer and 1 μL of Turbo DNAse (Thermo Scientific, cat***#***AM2238) were added, before a 30-min incubation at 37°C to eliminate genomic DNA. RNA was again purified by extraction with acid phenol:chloroform (Invitrogen, cat#AM9722) followed by precipitation with ethanol according to the manufacturer’s instructions, and then stored at -80°C or used immediately. 500 ng of RNA were used for reverse transcription with the SuperScript IV First-Strand Synthesis Kit (Invitrogen, cat#18091200) using random hexamers, according to the manufacturer’s instructions. RNA was degraded with 1 μL of RNase A (Thermo Scientific, cat# R1253) for 30 min at 37°C. The final product was diluted to 1:20 and 2 μL of the dilution were used in each 10-μl RT-qPCR reaction along with 400 nM of primers gene- or isoform-specific primers (Table S2B) and 1X SensiFAST SYBR No-ROX Mix (Bioline, cat#BIO-98005). Quantitative PCR was performed in CFX384 Touch Real-Time PCR Detection System (Bio-Rad). Each reaction was performed in triplicate. Normalized relative gene expression was determined using the mean Cq of each technical triplicate, and following the 2^-ΔΔCq^ method with *GAPDH* as a reference gene.

#### Purification of recombinant pore-forming toxins

PFT-expressing plasmids were purified from an overnight stationary culture of the corresponding strain by using the Wizard Plus SV Minipreps DNA purification System (Promega, cat#A1460) and transformed into *E. coli* BL21 (DE3) (New England Biolabs, cat#C2527H). Positive clones were screened by colony PCR, 3-4 positive colonies were inoculated in 20 mL of LB containing the appropriate antibiotic and incubated at 37°C, 180 rpm for 6 h to reach stationary phase. This starter culture was used to inoculate 1L of LB containing antibiotics. Incubation was pursued under shaking until a OD_600nm_ of 0.8 was reached. Isopropyl β-D-1-thiogalactopyranoside (IPTG) was then added at a final concentration of 0.5 mM and the culture was incubated overnight at 16°C. Bacteria were collected by centrifugation at 3, 600 × *g* for 10 min at 4°C. The pellet was washed once with ice-cold PBS and resuspended in lysis buffer (1.5 X PBS, 1 mM magnesium acetate, 0.1% IGEPAL, 20 mM leupeptin, 1 μg/mL pepstatin, 50 μg/mL PMSF and 1 X Lysozyme), sonicated for 4 min and centrifuged for 30 min at 27, 000 × *g*, 4°C. The supernatant was collected in a fresh 50 mL tube, mixed with 250 μL of Ni-NTA agarose resin (Invitrogen, cat#R901-15), that had been preequilibrated in an equal volume of lysis buffer, and the slurry was incubated for 2 h on a rotating device at 4°C. The resin was pelleted by centrifugation at 500 x g for 15 min, 4°C, washed twice with 5 mL of lysis buffer, then once with 500 μL of wash buffer (1.5X PBS, 250 mM NaCl, 1 mM magnesium acetate, 0.1% Igepal CA-630, 50 mM imidazole, 10% glycerol) and once more with 5 ml of lysis buffer. The beads, resuspended in 1 mL of lysis buffer, were transferred to a Poly-Prep Chromatography column (Biorad, cat# 731-1550) and the remaining supernatant was let to flow through. The slurry was incubated for 5 min on the column with 800 μL of elution buffer (1.5 X PBS, 1 mM magnesium acetate, 0.1% Igepal CA-3, 150 mM imidazole, 10% glycerol), then the eluate was collected, before proceeding to elution of another fraction (10 times). The concentration of each fraction was determined with Bradford Reagent (Abcam, cat#ab119216) and fractions with an absorbance above 0.1 were pooled and dialyzed overnight in 1L dialysis buffer (1.5X PBS, 1mM magnesium acetate, 10% glycerol and 2 mM dithiothreitol (DTT)) in 12-14 kDa Spectra/Por dialysis tubing (Repligen, cat#123697) with mild agitation at 4°C. The dialyzed protein solution was diluted in protein storage solution (20 mM MES pH 5.7, 150 mM NaCl, 5% glycerol, 2 mM DTT), aliquoted at 0.1 μM and stored at -80°C. Protein purity was assessed by running samples of every step of the purification process on SDS-PAGE and staining the gel using Quick Coomassie Stain (Neo-Biotech, cat#NB-45-00078) to reveal single bands in the eluted and dialyzed fractions.

Before use, pro-aerolysin was activated by incubation for 10 min with trypsin in a 1:20 aerolysin:trypsin molar ratio, then trypsin was inactivated by adding trypsin inhibitor (PromoCell cat#C-41120) to a final concentration of 0.02%.

#### Treatment of cells with pore-forming toxins or with pharmacological inhibitors

LoVo cells were seeded in 12-wells plates at an appropriate density to reach 85-90% confluence 48 h later. For TG003 treatment and PFTs treatment, the cells were washed once with PBS, then serum-free medium supplemented with PFTs at 4.5 nM, or with TG003 at a concentration ranging from 10 to 100 μM was added for 1h. Because aerolysin had been activated with trypsin, a similar concentration of inactivated trypsin diluted in the same buffer was added in the control sample. Cells were washed once with cold PBS, snap-frozen in liquid nitrogen and stored at −80°C until further use.

#### Ion fluxes perturbation

To test the role of calcium and potassium, isotonic calcium-free and high-potassium media were prepared as previously described (26). Potassium-free medium was prepared similarly to the Standard medium but lacked KCl. High-potassium^+^/calcium-free medium was prepared similarly to the high-potassium medium but lacked CaCl_2_. LoVo cells were seeded 48 h prior to experiment in a 12-wells plate format to reach 85-90% confluency. Cells were washed once with PBS and incubated for 30 min in the corresponding ion-altered medium prior to LLO 4.5 nM addition or not. Cells were incubated for another hour, then washed once with cold PBS, snap-frozen in liquid nitrogen and stored at −80°C until further use.

#### Long-read RNA-seq sample and library preparation

The RNA samples used for long-read cDNA sequencing (long-read RNA-seq) were the same samples that had been prepared in our previous work for total cytoplasmic RNA sequencing with Illumina next-generation sequencing technology (short-read RNA-seq) of *Lm*-infected LoVo cells over a 10-h time course (22). The integrity of isolated RNA was verified using the RNA Pico kit on an Agilent 2100 Bioanalyzer. Only high-quality RNA (RIN > 8) was used in library preparation.

Library preparation was performed using SQK-LSK108 following manufacturer’s protocol (1D PCR Barcoding cDNA; ONT) optimized for cDNA sequencing. Briefly, 100 ng of total RNA was reverse-transcribed for each sample with SuperScript IV (Life Technologies, cat#18090010), using custom polyT-VN and strand switching primers at 50°C for 10 min. Template switching was performed at 42°C for 10 min and enzyme inactivation at 80°C for 10 min. The reaction was purified with 0.7X Agencourt Ampure XP beads. A quarter of the purified RT product was taken into PCR for amplification and barcodes addition (Barcoded primers form EXP-PBC001, ONT), with a 17-min elongation at each 18 cycles. Double stranded cDNA was purified as above, quantified and checked for the size. Sample were multiplexed in equimolar quantities to obtain 1 μg of cDNA. The pool was end-repaired and dA-Tailed using the NEBNext End repair / dA-tailing Module (New England BioLabs, cat#E7546), and purified with 1X Agencourt beads. Adapter ligation was performed at 20°C for 10 min, with Adapter Mix (AMX, ONT) and Blunt/TA Ligase Master Mix (New England BioLabs, cat# M0367). After a final 1X clean-up and washing of the beads with Adapter Binding Buffer (ABB, ONT), the library was eluted in 15 μL Elution buffer. 900 ng of cDNA was loaded on an R9.4.1 flowcell after priming it according to the manufacturer’s protocol. Sequencing was performed with the standard 48-hour sequencing protocol run on the MinION MkIB, using the MinKNOW software (versions 1.7.1, 1.7.4, and 1.10.23). A mean of 1, 7 ± 0, 5 million passing ONT quality filter reads was obtained for each of the 12 samples. Base-calling from read event data was performed by guppy-3.6.1.

#### RNA Immunoprecipitation sample and sequencing library preparation (RIP-seq)

LoVo cells were cultured in 75 cm^2^ flasks and infected as described above. RIP was performed as previously described by Kwon et al. until RNA resuspension (27), except only immunoprecipitation (IP) 150 buffer was used for beads washing, and the antibodies used were anti-CIRBP or control IgG from rabbit serum (Sigma, cat#I5006). Two flasks per condition were pooled during final precipitation. Purified input or RIP-seq RNA samples were resuspended in 15 μL of water, their concentration was assessed by Qubit RNA High Sensitivity kit (Invitrogen, cat#Q32852) and samples were stored at -80°C until further processing.

For library preparation and Illumina sequencing, messenger (polyA^+^) RNAs were purified from 25 ng of RIP samples and 500 ng of total RNA input using oligo(dT). Libraries were prepared using the strand specific RNA-seq library preparation Stranded mRNA ligation kit (Illumina). Libraries were multiplexed by 12 for RIP samples and by 6 for input samples on P2 flowcell. A 118-bp single read sequencing was performed on a NextSeq 2000 (Illumina). A mean of 41 ± 19 million reads passing Illumina quality filter reads for RIP samples, and 88 ± 10 million reads for input samples, was obtained.

#### Read processing for long-read RNA-seq and RIP-seq

Fastq file quality was checked on Galaxy servers using the FastQC tool. Reads mapping to the 5S ribosomal RNA (OX177011.1) or 45S (NR_046235.3) pre-ribosomal RNA sequences with Bowtie2 (Galaxy Version 2.5.0) were filtered out. For long-read RNA-seq, filtered reads were mapped to the human genome (GRCh38.p13 from Gencode website) using minimap2 (Galaxy Version 2.24) with enabled spliced alignments. For RIP-Seq, filtered reads w ere mapped to GRCh38.p13 using RNA-STAR (Galaxy Version 2.7.8a) and a gene model file for splice junction (gencode.v41.annotation.gff3). Read processing for short-read RNA-seq data was described previously (22).

#### Alternative splicing analysis

Alternative splicing analysis was performed following the MAJIQ algorithm tutorial (28) using the IFB-core Cluster (https://www.france-bioinformatique.fr/en/ifb-core-cluster/). Alignment files for short-read RNA-seq were used along with Gencode annotation file gencode.v41.annotation.gff3 as inputs for the MAJIQ builder step. The deltaPSI method was used to run the MAJIQ quantifier, and results were modulized in VOILA (--changing-between-group-dpsi 0.05 --decomplexify-psi-threshold 0.01). Final results were visualized in the VOILA application as instructed. AS events were considered biologically relevant for |ΔPSI| between 10 hpi and non-infected cells over 0.1, and a probability of changing over 0.95. Functional enrichment analysis for genes significantly impacted by an AS event was conducted using over-representation analysis of GO biological processes with the gprofiler2 R package (version 0.2.1).

#### Long reads data analysis

Long-read RNA-seq alignment files were directly used to discover and quantify transcript isoforms by using the Bambu package in R (29) and the Gencode annotation file gencode.v41.annotation.gff3, generating a table of normalized transcript counts. From these tables, differential usage of the transcript isoforms was quantified with the IsoformSwitchAnalyzeR (ISAR) package in R (Vitting-Seerup and Sandelin, 2019, version 1.18.0). Information on impacted transcripts IDs and types were extracted from the Ensembl database (https://www.ensembl.org) using BioMart (version 2.52.0). Uniquely mapped reads were counted using FeatureCounts (version 2.0.1). Library size-normalized read alignments were visualized using the Integrative Genomics Viewer (IGV) from bedGraph files generated using samtools and bedtools.

#### RIP-Seq data analysis

Differential target enrichment of RIP-seq data were quantified by identifying enriched genes in CIRBP IP fractions compared (1) to the corresponding input fractions, and (2) to the corresponding control IgG IP fractions in two parallel analyses, using DESeq2 in R (version 1.36). CIRBP targets were considered as significantly enriched if they matched criteria of *p*_adj_ <0.05 and log_2_FC > 0.5 in both comparisons. For volcano plots, gene enrichment repartition for both non-infected and *Lm-* infected conditions was represented using values from the IP versus control IgG DESeq2 analysis. Functional enrichment analysis for significant CIRBP targets was conducted using over-representation analysis of GO biological processes with the gprofiler2 R package version 0.2.1.

#### Statistical analysis

Boxplots represent median and quartiles of data, with the mean plotted as a diamond shaped point and each individual value as a full black dot. Information on independent biological replicates and about the statistical tests used is provided in the caption for each graph. For immunoblots, protein band intensities were quantified using the Fiji software, and normalized to the corresponding housekeeping protein. Mean intensity values were compared using Student’s *t*-test.

## Results

### Infection of epithelial cells by *Listeria monocytogenes* alters the mRNA alternative splicing landscape

While quantitative aspects of human gene expression changes upon *Lm* infection have been analyzed in-depth in a number of transcriptomic and translatomic studies (22, 31), we sought to explore whether gene expression was also affected at the mRNA isoform level, which could impact the function of protein products. To this end, we investigated AS events affecting human transcripts at several time points (0, 2, 5, and 10 h) post-infection (hpi) with *Lm* LL195 strain in the LoVo epithelial cell line of colon adenocarcinoma. We first re-analyzed short-read RNA sequencing (RNA-seq) data that we had previously generated across the 10-h infection time-course (22) using the MAJIQ algorithm, a method optimized for the identification of local splicing variations (LSVs) regardless of isoform-based alignments (28). The high-depth of Illumina sequencing of cDNA fragments allowed a robust detection of statistically-significant changes of splice sites usage across conditions (Tables S3A to S3C), while long-read cDNA sequencing of the same set of samples obtained with MinION (Oxford Nanopore Technologies) proved better adapted for qualitative investigation of isoform switches in a selected subset of candidate genes at later stages of investigation, but lacked depth for *bona fide* statistical analysis of isoform switches (Tables S3D to S3F). Relative changes in AS occurrence from one condition to another were quantified as the difference in percent spliced-in (PSI), dubbed delta PSI (dPSI) (28). Whereas almost no change in AS events were detected at 2 hpi and 5 hpi when compared to non-infected (NI) cells (4 and 5 LSVs with over 5% dPSI and 95% probability changing, respectively), a defined subset of LSVs (417), spanning over 191 genes, were significantly impacted by *Lm* infection at 10 hpi, with no overall gradual pattern across time-points (Figure 1A and 1B). Gene ontology overrepresentation analysis of genes affected by LSVs at 10 hpi revealed a significant enrichment in functions associated with RNA metabolism, and notably with splicing itself (Figure 1C). Alteration of splicing in transcripts from the SRSF family (Figure 1A, Figures S1A and S1B) is consistent with well characterized auto-regulatory feedback loops that govern the expression of genes encoding SR splicing factors (32–34). Together, these results suggest that AS-based regulations contribute to the fine-tuning of the expression of a subset of human genes in response to *Lm* infection, some of which can themselves have a role in downstream host gene expression regulation. We then sought to investigate in more detail the alternative isoforms of one of these candidate regulators, the upstream pathways leading to their processing, and their potential downstream role in infection.

**Figure 1.**
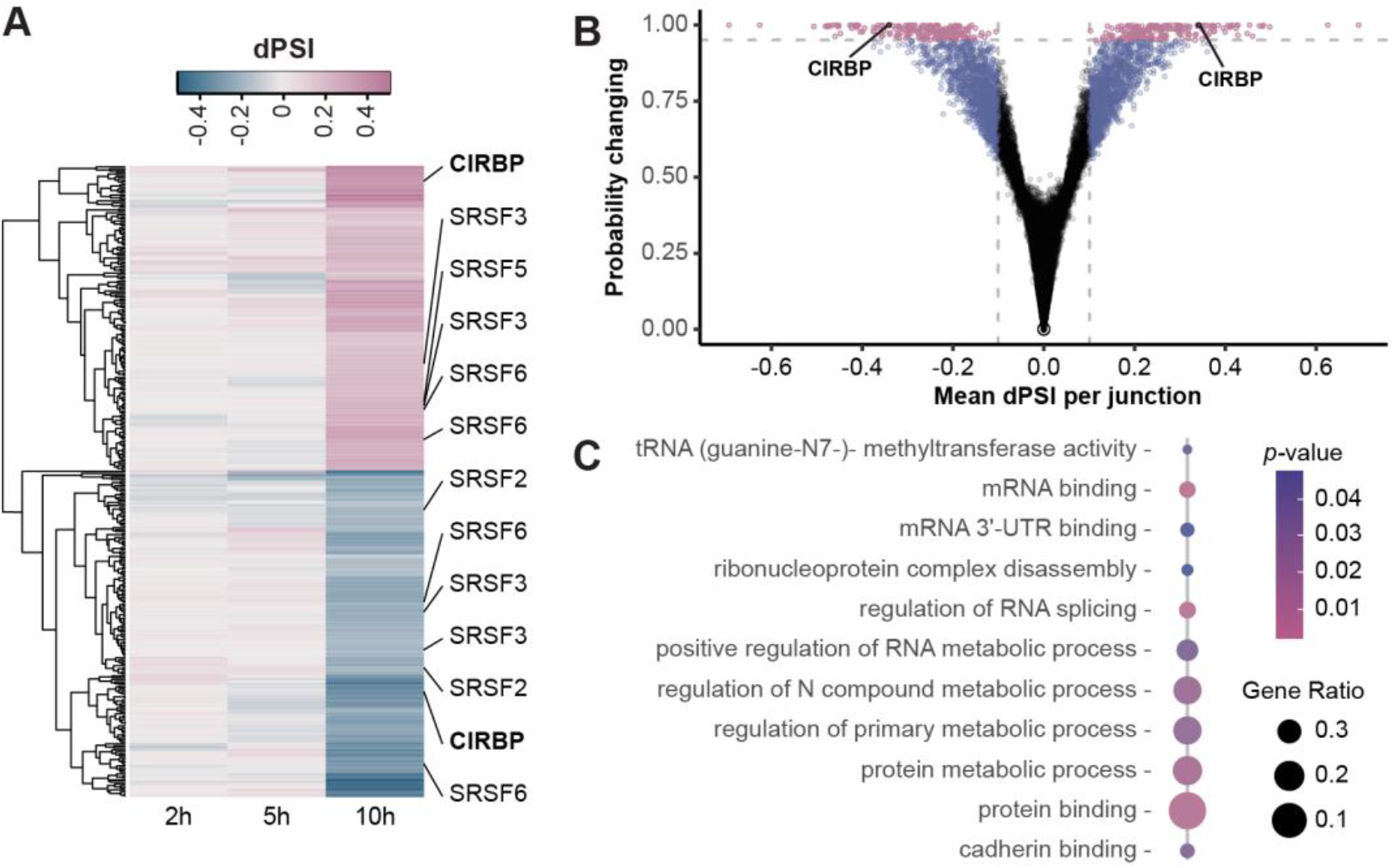
*Listeria monocytogenes* infection alters the representation of mRNA isoforms for a subset of human genes, including CIRBP. (A-B) Variations in RNA local splicing variants in LoVo intestinal epithelial cells infected for 2, 5 or 10 h with *Lm*. Short-read cDNA sequencing data were used for the detection of LSVs using the MAJIQ algorithm (3 independent replicates at 0, 2, 5 hpi; 2 at 10 hpi). (A) Heatmap of the difference at 2, 5 and 10 hpi in percentage spliced-in (dPSI) values of significant AS events induced by *Lm* infection, compared with non-infected cells (probability of changing > 0.95). The names of a subset of genes positively (pink) and/or negatively(blue) affected by LSVs are indicated. (B) Volcano plots highlighting the most significant AS events in *Lm*-infected cells at 10 hpi compared to non-infected cells. Vertical dashed grey lines indicate a mean dPSI of 0.1 and the horizontal dashed gray line indicates a probability of changing of 0.95. (C) Over-representation analysis of gene ontology biological process terms for genes undergoing RNA isoform-based regulation in LoVo cells at 10 hpi compared to non-infected conditions (probability of changing > 0.95; all dPSI beyond this threshold were above 0.1).

### CIRBP is alternatively spliced at the junction of exons 6 with downstream exons

Among the genes affected by the most robust isoform switches, CIRBP (cold-inducible RNA binding protein) struck us as worthy of functional investigation. Indeed, the reported roles of CIRBP isoform-based regulation in orchestrating cellular responses to different types of cellular damages prompted us to further document its status upon *Lm* infection. High dPSI and probability of changing for two mutually exclusive LSVs detected at the junction of exon 6 with either 7a or 7b by short-read analysis (Figure 1A and B, Table S3C) was matched by high up-*versus* down-regulation of isoforms mutually differing in the alternative use of exons 7a versus 7b in long-read RNA-seq analysis (Table S3F, Figure 2A and B). By contrast, no statistically significant LSV affecting the 5 first exons of CIRBP was detected, prompting us to focus thereafter on isoforms that varied in their distal part. More specifically, the junction between a splicing donor site in exon 6 located at position 1, 272, 051 in the genome and downstream acceptor sites was the only statistically significant LSV between infected and non-infected conditions. Among the 46 splice variants that are currently annotated for human CIRBP, we grouped those that were detected in our analysis into three main classes of isoforms differing at the junction of exon 6 (Figure S2): (*a*) a dominant protein-coding class bridging exon 6 with exon 7a, the most representative of which is <u>ENST00000320936.9</u>, CIRBP-201, that accounted for 89.6±12.0% of CIRBP isoforms in non-infected cells (Table S3G, long-read data); (*b*) a class of alternatively spliced transcripts bridging exon 6 with exons 7b the prototype of which being the predicted NMD target <u>ENST00000586636.5</u>, CIRBP-210 (9.8±1.9% of transcripts in non-infected cells; Table S3G, long-read data); and (*c*) a third class encoding elongated versions of the CIRBP protein due to the retaining of introns with coding potential after exon 6, best represented by CIRBP-225 (<u>ENST00000589710.5</u>) (0.6±0.8% of transcripts in long-read data in non-infected cells; Table S3G). By 10 hpi, CIRPB-210 expression underwent a 2.6-fold reduction in profit of a 9.5-fold increase for CIRBP-210-like isoforms, which then represented 71.9±19.8% of CIRBP transcripts (Figure 2A and B, Table S3G, long-read data). Due to extremely low counts, variations in CIRBP-225-like isoforms at 10 hpi were irrelevant (Table S3G), and no change in CIRBP-225 expression was detected by RT-qPCR using primers specific for the retained intron in this isoform (Figure S3). Subsequently, we focused on the two most represented isoform classes (*a*) and (*b*) affected by a statistically significant LSV.

**Figure 2.**
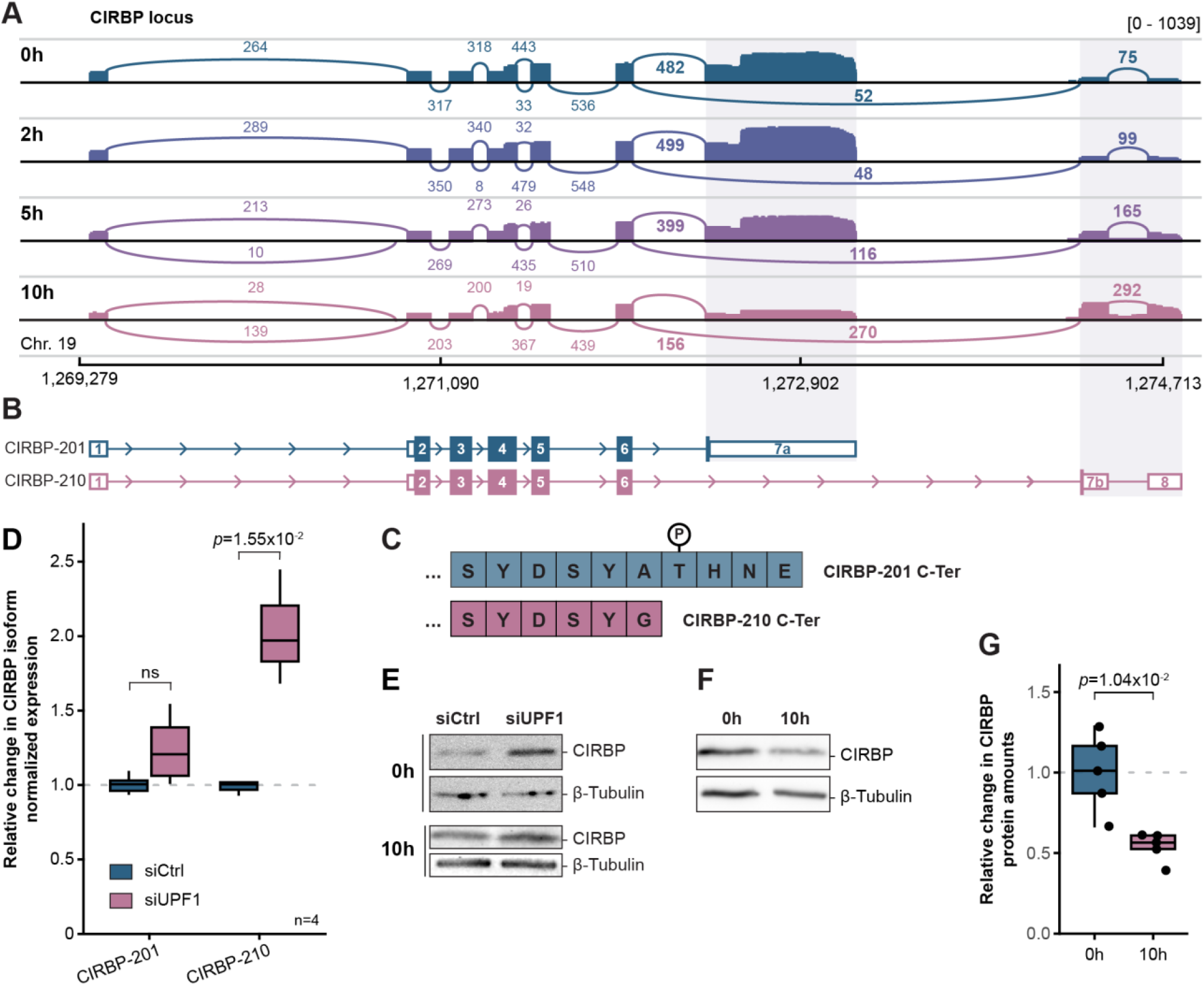
Alternative splicing of CIRBP leads to a switch towards an NMD-sensitive isoform upon *Listeria monocytogenes* infection. (A) Sashimi plot of long-read Nanopore sequencing data aligned at the CIRBP locus. For each timepoint, average values of read counts per genomic position normalized for library size, and exon-junction coverages for each junction, are represented (3 independent replicates at 0, 2, 5 hpi; 2 at 10 hpi). The range for normalized read counts is indicated between brackets. (B) Representation of the mature forms of the two prototypical transcripts of CIRBP, CIRBP-201 and CIRBP-210, aligned with their position on the genome in (A). Exons are represented as colored boxes, the height of which is magnified in the CDS; introns are displayed as lines with arrows indicating 5’- to 3’-orientation. Alternative 3’-ends, corresponding to exons 7a in CIRBP-201, and 7b-8 in CIRBP-210, are shaded in gray. (C) Last amino acids in the sequence of the proteins encoded by CIRBP-201 (top, in blue) and CIRBP-210 (bottom, in pink). The circled P indicates the site of a putative threonine phosphorylation site. (D Relative CIRBP-210 to -201 transcript ratio in LoVo cells transfected with a siRNA targeting UPF1 (siUPF1) or a scrambled siRNA (siCtrl). For each condition, the quantities of CIRBP-210 and CIRBP-201 isoforms normalized to GAPDH were quantified by RT-qPCR, and normalized to their expression levels in cells treated with a scrambled siRNA (siCtrl). Boxplots represent medians and quartiles of data from four independent experiments. One way ANOVA followed by post-hoc Tukey’s test was used for statistical testing between conditions. (E) Assessment of CIRBP protein abundance by immunoblot on lysates from LoVo cells transfected with a siRNA targeting UPF1 (siUPF1) or a scrambled siRNA (siCtrl), either uninfected (0 h) or infected with *Lm* for 10 h. (F) Assessment of CIRBP protein abundance by immunoblot on lysates from LoVo cells infected for 10 h with *Lm*, compared to non-infected cells (0 h). (G) CIRBP protein quantification on 5 independent experiments performed as in (F). On each immunoblot, mean CIRBP intensities were normalized to mean intensities for tubulin. Student’s *t*-test was used for statistical testing between conditions.

### Alternative use of 3’ exons affects CIRBP mRNA expression in infected cells

CIRBP-201 and -210 differ in their 3’ regions, which consist of alternative exons 7a versus 7b-8 (Figure 2B). The impact of the AS on the CDS is minimal, giving rise to a 4-amino acid shorter protein for CIRBP-210 compared to CIRBP-201, and to the loss of a putative phosphorylation site on a threonine (35) (Figure 2C). By contrast, their 3’-untranslated regions (UTRs) differ significantly (Figure 2B). Indeed, in CIRBP-210, the presence of an exon-exon junction in the 3’-UTR makes it a target for degradation by NMD (9), which is attested by its sensitivity to the depletion of key NMD factors (data from 36). In the LoVo cell line, we confirmed that RNA silencing of the central NMD factor UPF1 resulted in increased CIRBP-210 mRNA abundance and CIRBP protein levels compared with non-transfected cells (Figure 2D and E), indicating that NMD destabilizes CIRBP-210-like isoforms.

Upon infection, no general upregulation of annotated human NMD-sensitive transcripts was observed in long-read RNA-seq datasets (Figure S4), ruling out that the increased abundance of CIRBP-210 was due to the impairment of central NMD pathways. This result also implies that, although the production of CIRBP isoforms was shifted in favor of CIRBP-210 in infected cells, this class of transcripts was still sensitive to NMD, and thus less stable than its -201 counterpart. In agreement with CIRBP-210 being less stable than CIRBP-201, the overall mRNA abundance of CIRBP was decreased 1.6-fold at 10 hpi (Figure S1B). Its coverage in ribosome footprints was reduced 2.8-fold, corresponding to a 1.7-fold drop in translation efficiency.

Immunoblotting against CIRBP confirmed that down-regulation of its expression was part of the consequences of *Lm* infection (Figure 2F and G). However, CIRBP was previously reported to have a half-life over 8 h in four different cell lines (37). It is thus unlikely that the down-regulation of CIRBP mRNA levels is sufficient for the observed reduction in CIRBP protein abundance at 10 hpi (Figure 2F). On top of RNA-based regulation, the loss of intracellular CIRBP may thus either be driven by an acceleration of CIRBP turnover rate, or by its secretion under the form of eCIRBP, as previously reported to occur in cell damaging conditions (38). Indeed, while UPF1 silencing resulted in a rise of CIRBP-210 mRNA abundance at 10 hpi comparable to its effect in uninfected cells (Figure S3), CIRBP protein levels were only modestly increased (Figure 2E), suggesting that infection-induced CIRBP degradation or secretion dampens the up-regulation of CIRBP protein by UPF1 silencing.

### Infection-induced CIRBP isoform switch requires LLO-proficient *Lm* strains

To investigate whether specific virulence properties of *Lm* were mediating the switch in CIRBP isoform usage upon infection, we compared the ability of distinct *Listeria* strains to alter the proportion of CIRBP-201 and -210 isoforms in LoVo cells. Each isoform class was quantified by RT-qPCR using primers specific to alternative terminal exons 7a or 7b-8, then the ratio of CIRBP-210 to -201 relative abundance, normalized to its value in non-infected cells, was compared across conditions (Figure 3A). For wild-type (WT) *Lm*, while no significant change was observed at 5 hpi, CIRBP-210 to -201 abundance was increased 1.6-fold at 10 hpi, confirming CIRBP isoform switch. By contrast, neither a 10-h infection with a strain of *Listeria innocua* rendered invasive by carrying *Lm* internalin gene *inlA* (*L. innocua* [*inlA*]), nor with a strain of *Lm* that is deficient for the *hlyA* virulence gene, encoding the pore-forming toxin (PFT) listeriolysin O (*Lm* Δ*hlyA*), affected CIRPB-210 to -201 relative abundances. Consistently, CIRBP protein levels were reduced in cells infected for 10 h with WT *Lm*, while they were not in cells infected with *Lm* Δ*hlyA* (Figure S5A). As *L. innocua* does not harbor the *hlyA* gene, these results strongly suggest that LLO is a key determinant of CIRBP isoform switch, which is coherent with the activity of LLO being well-known to activate cellular stress response pathways (22–25).

**Figure 3.**
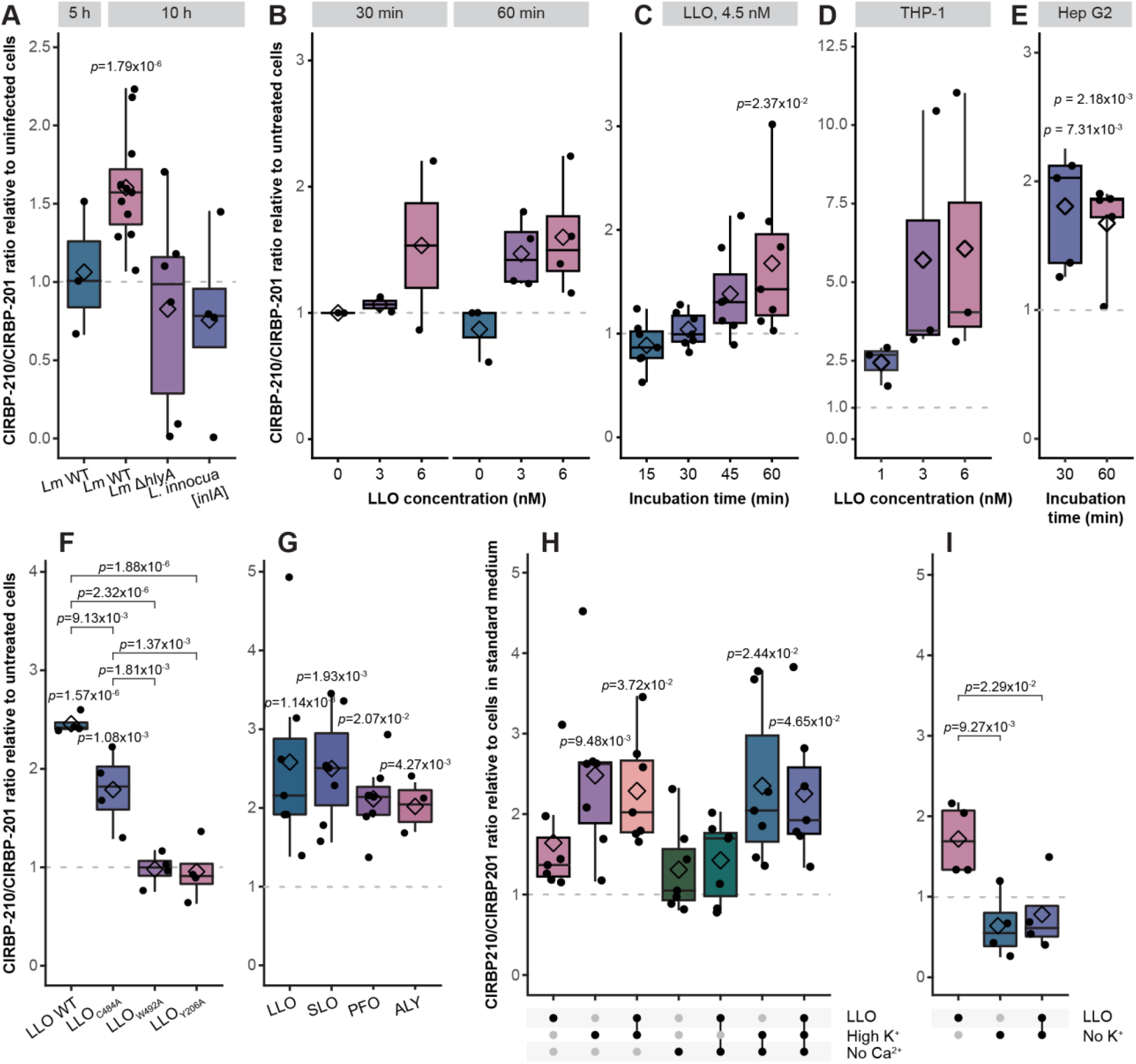
Bacterial pore-forming toxins induce a change in the balance between CIRBP isoforms. (A-I) Quantification of the relative abundance between CIRBP-210 and -201 transcripts in cells in response to infection or to exposure to bacterial PFTs. For each condition, the ratio of CIRBP-210 to CIRBP-201 isoforms was quantified by RT-qPCR, and normalized to the CIRBP-210/CIRBP-201 ratio in uninfected, untreated cell (dashed grey lines). Boxplots represent medians and quartiles of data, with the mean represented as diamonds. Two-way ANOVA followed by post-hoc Tukey’s test was used for statistical testing between conditions. Each dot represents an independent experiment. (A) Relative CIRBP transcript ratio in LoVo cells infected for 5 h with wild type (WT) *Lm*, or for 10 h with WT *Lm*, with *Lm* deleted for the *hly*A gene (Δ*hlyA*), or with a strain of *L. innocua* rendered invasive by the expression of *Lm* internalin gene *inlA*, compared to uninfected LoVo cells. (B) Relative CIRBP transcript ratio in LoVo cells treated for 30 or 60 min with 3 or 6 nM of listeriolysin (LLO), compared to untreated cells. (C) Relative CIRBP transcript ratio in LoVo cells treated for 15 -to 60 min with 4.5 nM of LLO compared to untreated cells. (D) Relative CIRBP transcript ratio in PMA-activated THP-1 monocytes treated for 1 h with 1, 3 or 6 nM of LLO, compared to untreated cells. (E) Relative CIRBP transcript ratio in Hep G2 hepatocytes treated for 30 or 60 min with 4.5 nM of LLO compared to untreated cells. (F) Relative CIRBP transcript ratio in LoVo cells treated for 1 h with 4.5 nM of LLO or of catalytically-impaired mutants of LLO, compared to untreated cells. (G) Relative CIRBP transcript ratio in LoVo cells treated for 1 h with 4.5 nm of CDCs (perfringolysin O, PFO; streptolysin O, SLO; or aerolysin, ALY), compared to untreated cells. (H and I) Relative CIRBP transcript expression ratio in cells incubated in isotonic buffers containing varying cation concentrations 30 min, before addition or not of 4.5 nM of LLO for 1 h. (H) Incubation in standard medium containing 140 mM NaCl, 5 mM KCl, 1.5 mM CaCl_2_, compared with high potassium medium containing 140 mM KCl, 5 mM NaCl, 1.5 mM CaCl_2_ (High K^+^) or calcium-free medium (No Ca^2+^). (I) Incubation in standard medium, compared with medium lacking extracellular potassium (No K^+^).

### The pore-forming activity of LLO is sufficient to induce CIRBP isoform switch

We next sought to determine whether the pore-forming activity of LLO was mediating the switch in CIRBP isoform usage. Incubation of LoVo cells with 3 to 6 nM of purified LLO for 15 to 60 min shifted CIRBP AS in favor of CIRBP-210 in a dose- and time-dependent manner (Figure 3B and C). This phenotype was not specific to the LoVo epithelial adenocarcinoma cell line, as it was reproduced in PMA-activated THP-1 macrophage-like cells as well as in Hep G2 hepatocytes (Figure 3D and E).

CIRBP protein levels were also decreased upon treatment with increasing concentrations of LLO (Figure S5B), arguing that the CIRBP loss observed during infection is LLO-dependent (Figure 2F and G, Figure S5A). In addition, LLO treatment for 30 to 60 min resulted in increased amounts of eCIRBP detected in the supernatants of both LoVo and THP-1 cells (Figure S5C). This observation suggests that secretion of CIRBP in the extracellular environment contributes to the decrease of intracellular CIRBP protein upon exposure to LLO, as it was hypothesized above during infection.

We then incubated LoVo cells with 4.5 nM of purified WT or mutated versions of LLO for 1 h, before assessing the relative abundance of CIRBP isoforms (Figure 3F). Two of the variants of LLO used are mutated in the conserved undecapeptide of cholesterol-dependent PFTs (C484A or W492A), resulting in a respective loss of 70% and over 99% of their hemolytic activity; the third mutant (Y206A) also displays an over 90% loss of hemolytic activity (39). Exposure to WT LLO induced a 2.5-fold increase in the CIRPB-210 to -201 ratio, and the LLO mutant retaining 30% of its hemolytic activity also induced a 2-fold isoform switch. By contrast, catalytically inactive LLO mutants had no effect on CIRBP transcripts, thus emphasizing that LLO was sufficient to stimulate CIRBP isoform switch, and that its pore-forming activity was mechanistically required in this process.

### The switch in CIRBP isoform can be induced by a broad array of pore-forming toxins

As mentioned above, LLO belongs to the extended family of PFTs; more specifically, it is a cholesterol-dependent cytolysin (CDC) that creates 30-nm pores in membranes, although it can also be inserted as arcs or incomplete rings (40). To clarify the properties of cell membrane permeation that were responsible for LLO-mediated switch in CIRBP isoforms, the ability of other purified PFTs to induce this switch was assessed. Similarly to LLO, streptolysin O (SLO) and perfringolysin O (PFO) are CDCs that create large pores, while aerolysin (ALY) does not bind cholesterol but GPI-anchored proteins and forms pores of smaller diameter (1.5-2 nm) (23, 41). Using identical incubation conditions as for LLO —except that pro-aerolysin was activated for 10 min with trypsin (42)—, the three PFTs induced a 2.2- to 2.6-fold increase in isoform ratio (Figure 3G). CIRBP isoform switch can thus be driven by the perforation of membranes, independently on pore-size. This suggests that a CIRBP-regulating cellular stimulus could be induced by the transit of ions or small solutes across the plasma membrane.

### Perturbations of cationic gradients across the plasma membrane phenocopies LLO-induced switch in CIRBP isoform abundance

By forming pores in the plasma membrane, LLO and other PFTs lead to potassium effluxes as well as calcium and sodium influxes, thus altering the gradients of cation concentrations that are actively maintained across the plasma membrane in living cells (26, 43). To test if LLO-dependent cation leakage was involved in CIRBP regulation, cells were incubated in conditions that inhibit passive ion fluxes by using a high-potassium/low sodium medium, to prevent potassium efflux/sodium influx, a calcium-free medium, to prevent calcium influx or a high-potassium and calcium-free medium to prevent both fluxes. Sodium concentrations were adjusted so that each medium remained isotonic, in order to avoid osmotic shock. Cells were pre-incubated for 30 min in each medium prior to LLO addition for 1 h, then CIRBP isoform ratio was assessed by RT-qPCR (Figure 3H). While cells incubated in the calcium-depleted medium behaved like cells in standard medium, with a 2-fold increase in the CIRBP-210 to -201 ratio in presence of LLO, a high extracellular potassium/low sodium concentration alone was sufficient to induce a stronger switch of CIRBP isoforms than LLO, whether in presence or absence of LLO or in calcium-depleted medium. Conversely, totally eliminating potassium from the extracellular medium led to a decrease in CIRBP-210/CIRBP-201 ratio that was not compensated by incubation with LLO (Figure 3I), suggesting that the balance between sodium and potassium concentrations across the plasma membrane has a strong influence on CIRBP regulation. Since LLO provokes potassium effluxes through membrane pores, one possible hypothesis for the nature of the signal that triggers CIRBP isoform switch would be a sensing of extracellular potassium concentration increase (and/or sodium decrease), or a cytotoxic effect of this ionic imbalance. An alternative —although intimately related— hypothesis would be that the CIRBP isoform switch is triggered by perturbations of the plasma membrane potential as a result of cationic imbalance across the plasma membrane.

### CIRBP alternative splicing is under control of CLK1 kinase activity in LoVo cells

A switch in CIRBP alternatively-spliced isoforms, similar to the one we observed in response to *Lm* infection and to the activity of PFTs, has already been reported to occur as a consequence of heat stress (7, 9). This regulation has been directly linked to a loss of activity of the CLK1 kinase (7). Upon heat stress, CLK1 inhibition has been shown to shift the splicing of CIRBP in favor of CIRBP-210-like isoforms, likely through the dephosphorylation of serine/arginine-rich splicing factors, known as SR proteins, that are targets of CLK1 kinase activity (7, 9). Consistently, treatment of LoVo cells with 0 to 5 μM of the CLK1 pharmacological inhibitor TG003 for 48 h, or with 75 μM of TG003 for 1 h, shifted the splicing of CIRBP pre-mRNAs in favor of CIRBP-210 in a dose-dependent manner (Figure 4A), thus validating that CLK1 is a regulator of CIRBP isoform switch in LoVo cells, similar to what has been described in the HEK293 cell line (7).

**Figure 4.**
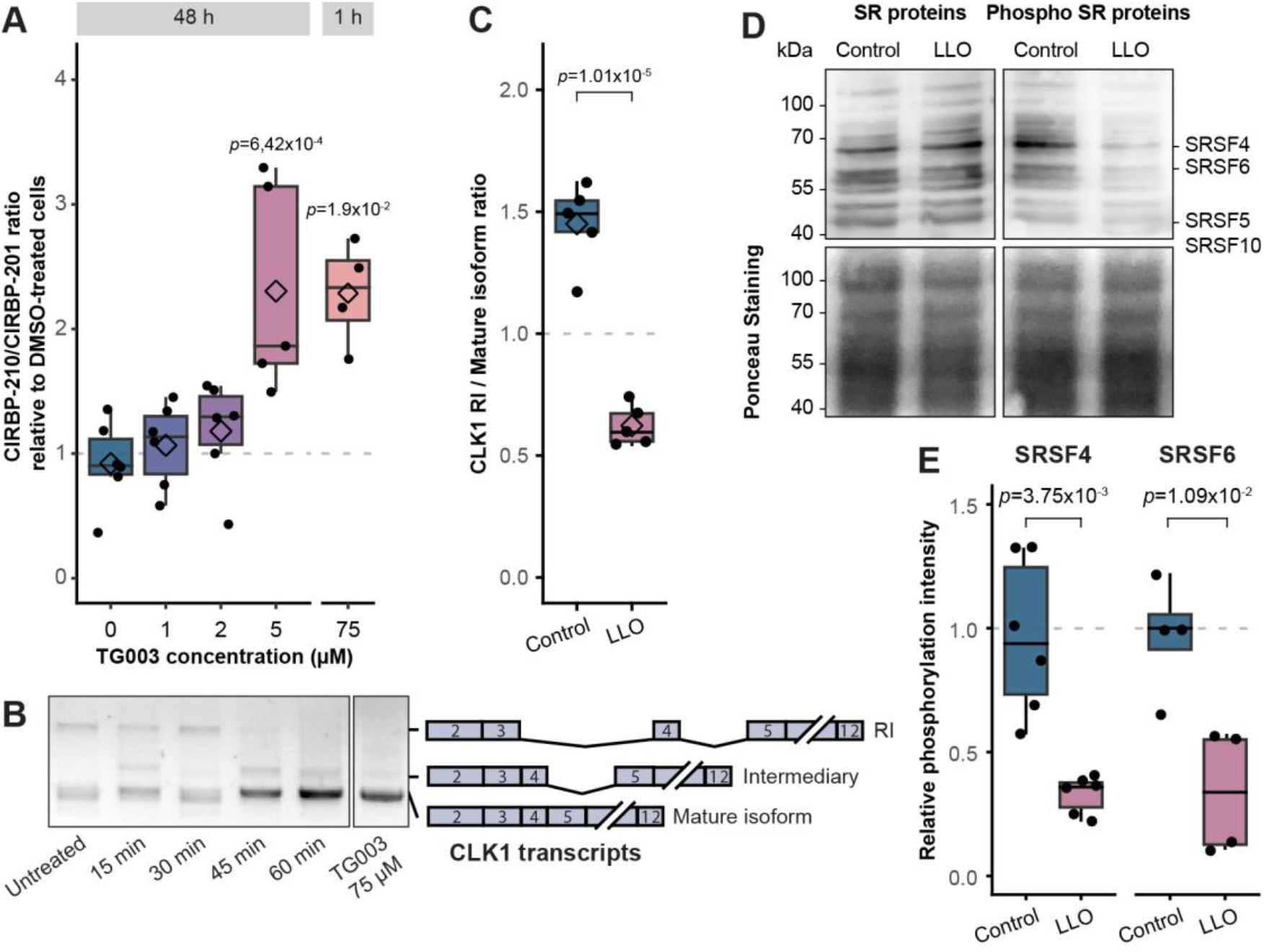
CLK1 activity is regulated in response to LLO-induced stress and impacts CIRBP alternative splicing in LoVo cells. (A) Relative CIRBP transcript ratio in cells incubated for 48 h with 0.5 to 5 μM of the CLK1 inhibitor TG003 (n = 5), or for 1 h with 75 μM of TG003 (n = 4). RT-qPCR quantification was performed and normalized as detailed in the legend to Figure 3. (B) Assessment of the splicing of CLK1 retained introns 3 and 4 upon incubation with LLO. LoVo cells were treated with 4.5 nM of LLO for 15 min to 1 h, or with 100 μM of TG003 (a pharmaceutical inhibitor of CLK1) for 1 h, then the maturation of CLK1 transcripts was assessed by reverse transcription followed by PCR with specific primers. (C) Quantification by RT-qPCR of the proportion of the retained-intron (RI) isoform with respects to the mature isoform in cells treated for 60 min with 4.5 nM of LLO compared to untreated cells in five independent experiments. For each condition, the abundance of each isoform was quantified with two pairs of specific primers spanning relevant exon-exon or exon-intron junctions and normalized to GAPDH. The average of the two pairs was used for calculating the ratio between the RI and mature isoforms. (D) Assessment by immunoblotting of the phosphorylation status of SRSF proteins in LoVo cells incubated for 1 h with LLO compared to untreated cells (representative experiment from 4 replicates). (E) Quantification of the phosphorylation of SRSF4 and SRSF6 from, respectively, 4 and 6 independent immunoblots performed as in (C). For each condition, the mean intensity detected with the anti-phospho-SR protein antibody was normalized to the corresponding intensity for GAPDH.

### The processing of CLK1 pre-mRNA is altered upon infection

The expression of CLK1 itself is highly sensitive to heat stress (44). Indeed, CLK1 transcripts are represented by two major forms: a mature protein-coding mRNA, and a premature, nuclear form in which introns 3 and 4 are retained (Retained Introns, RI). Upon heat stress, the dephosphorylation of CLK1 prevents its kinase activity on SR splicing factors (7, 44). The subsequent dephosphorylation of SR proteins promotes the splicing of the RI pool into mature CLK1 mRNAs, which facilitates the rapid re-phosphorylation of SR splicing factors upon stress release (44). To assess the effect of LLO-induced stress on CLK1 maturation, a region spanning exons 2 to 12 was amplified by PCR on cDNA prepared from cells that had been exposed to 4.5 nM LLO for 15, 30, 45, or 60 min, or to 75 μM of TG003 for 1 h, or from untreated cells (Figure 4B). The pore-forming toxin stimulated the processing of CLK1 pre-mRNA from the RI form (top band) into its mature form (bottom band), with an intermediate form where the splicing between exons 3 and 4 was restored appearing after 15 min of treatment. The fully-spliced transcript accumulated after 45 min, until being nearly fully-spliced after 1 h of treatment, phenocopying the effect of 1 h of TG003-mediated CLK1 inhibition. Quantification by RT-qPCR of the ratio between the RI and the mature isoform confirmed the loss of the RI form after 60 min of treatment with 4.5 nM of LLO compared to untreated cells (Figure 4C). LLO treatment can thus recapitulate the auto-regulatory processing of CLK1 transcripts that is a typical signature of CLK1 loss of activity and SR protein dephosphorylation, similar to what occurs upon heat stress (44).

### The phosphorylation and expression of SR splicing factors are impaired upon infection or LLO incubation

To determine the impact of *Lm*- or LLO-mediated regulation of CLK1 on its activity, the phosphorylation status of SR proteins was detected by specific immunoblotting against their phosphoepitopes at 1 h post-treatment of cells with 4.5 nM of purified LLO. LLO induced a reduction of the phosphorylation level of SR proteins, best represented here by SRSF4 and SRSF6, by about 70% (Figure 4D and 4E).

The effects of SR proteins dephosphorylation on their activity is expected to affect the splicing of other genes than CIRBP, including SRSF genes that encode SR proteins themselves (32, 34, 45). Consistently, nine SRSFs were affected by AS events at 10 hpi compared to NI cells (Figure 1A, Figures S1A and S6). Among these, five (SRSF2, 3, 4, 6, and 7) were affected by shifts towards transcripts that harbor a “poison exon” rendering them targets for degradation by NMD, and three (SRSF1, 5 and 10) resulted in noncoding alternative transcripts. Accordingly, the expression of these genes was dampened in infected cells, as attested by lower RNA levels and ribosome footprints (Figure S1B). These results corroborate a model in which the stress generated during infection by the pore-forming activity of LLO leads to the dephosphorylation of SR-proteins *via* CLK1 inhibition, which in turns reshapes the cellular splicing landscape, including by favoring the splicing of CIRBP pre-mRNAs towards NMD-sensitive isoforms, as well as that of SRSF pre-mRNAs towards unstable or noncoding transcripts.

### CIRBP isoforms have opposite effects on cell infection

To investigate the possible consequences of CIRBP regulation as a player in the *Lm*-host crosstalk, isoform-specific knockdown was performed using siRNAs targeting (*a*) the CDS of CIRBP, which targets both CIRBP-201 and -210 (siCIRBP-CDS), (*b*) the 3’-UTR of CIRBP-201 specifically (siCIRBP-201), or (*c*) the 3’-UTR of CIRBP-210 specifically (siCIRBP-210) (Figure S7). After 48 h of RNA interference, cells were infected with *Lm* for 10 h before enumerating intracellular bacteria (Figure 5A). Knocking down CIRBP-210 led to a 27% increase in colony forming units (cfu) per cell, while knocking down CIRBP-201 decreased the number of cfu/cell by 31%. Knock-down of all CDS-containing CIRBP isoforms displayed an intermediate phenotype with a non-significant 14% decrease in infection. These results hint at a role for CIRBP in the control of cell autonomous responses to infection, and emphasize the contribution of CIRBP isoform switch in opposing the colonization of cells by bacteria by dampening the expression of a pro-bacterial protein-coding isoform in favor of a less stable transcript.

**Figure 5.**
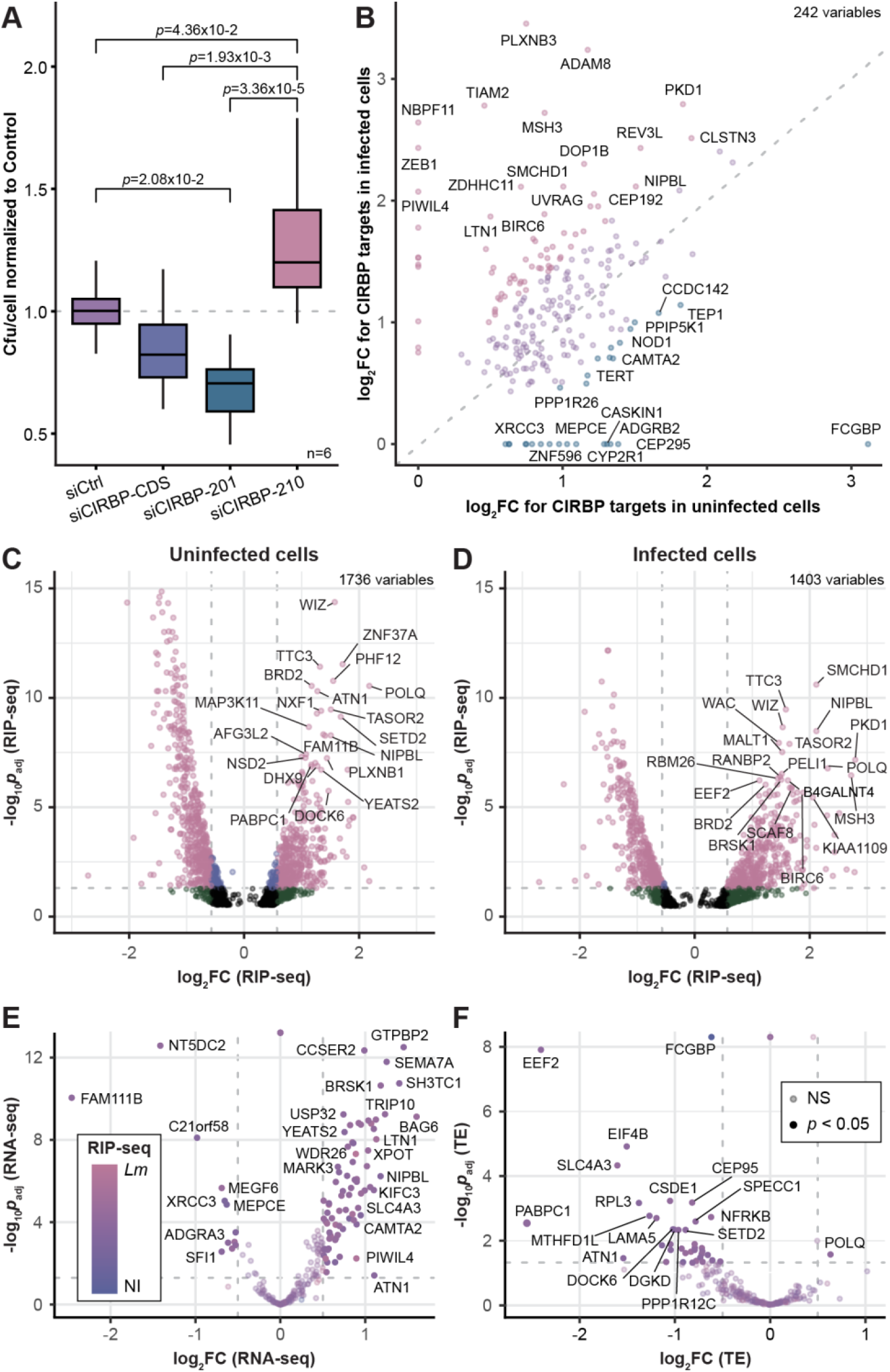
Function of CIRBP in the cellular response to *Listeria monocytogenes* infection. (A) Quantification of *Lm* infection in LoVo cells silenced for CIRBP, or for one of its isoforms. Bacterial colonisation of LoVo cells, transfected with siRNA targeting the CIRBP CDS (siCIRBP-CDS), CIRBP-201 3’-UTR, CIRBP-210 3’-UTR or a control scrambled siRNA (siCtrl), was assessed by gentamicin protection assay followed by serial dilution plating of infected cell lysates on agar plates. The number of colony-forming unit (cfu) per cell in each condition was normalized to infection levels in the control condition in 6 independent experiments. One-way ANOVA followed by post-hoc Tukey’s test was used for statistical testing between conditions. (B-D) RNA-immunoprecipitation of CIRBP targets. CIRBP targets were identified by differential expression analysis between a CIRBP-pull down mRNA fraction and both total mRNAs (input) or a control IgG pull-down mRNA fraction. Values of enrichments between the CIRBP pull-down fraction and the IgG pull-down fraction were used for plotting. Pseudogenes were not plotted. Data from two independent replicates. FC, fold change; *p*_adj_, adjusted *p*-value [DESeq false discovery rate (FDR)]. (B) Scatter plots of changes in enrichment of significant CIRBP targets in uninfected *versus Lm*-infected cells. Only transcripts enriched in the CIRBP pull-down fraction in both comparisons (against the input, and against control IgGs) were kept as targets (2, 542genes). Data points colored in mauve represent transcripts that were similarly bound by CIRBP in the two datasets (|Δlog_2_FC*_Lm_*_-NI_| < 0.5), while points colored in pink (Δlog_2_FC > 0.5) and blue (Δlog_2_FC < -0.5) correspond respectively to transcripts enriched in infected and uninfected cells. (C-D) Volcano plots highlighting CIRBP targets being significantly enriched in (C) uninfected cells and (D) *Lm-*infected cells at 10 hpi compared to the control RIP. Data points colored in pink represent genes with *p*_adj_ below 0.05 (dashed grey horizontal line, -log_10_ *p*_adj_ = 1.3) and a FC above 1.41 (vertical dashed grey lines, log_2_ FC = ± 0.5). Grey dots, non-significant; green dots, log_2_FC > 0.5, *p*_adj_ > 0.05; blue dots, *p*_adj_ < 0.05, log_2_FC < 0.5; pink dots, log_2_FC > 0.5 and a *p*_adj_ < 0.05. The 20 genes with the highest Manhattan distance to the origin were labeled on each graph. (E-F) Volcano plots highlighting changes in mRNA levels (RNA-seq data; E) and translation efficiency (TE) assessed by normalizing Ribo-seq counts to RNA-seq counts (F) for the 242 CIRBP targets, in cells infected with *Lm* for 10 h compared to non-infected cells. The color of data points ranging from blue to pink reflects the Δlog_2_FC of RIP enrichment in infected (*Lm*) versus non-infected (NI) cells. Data points colored in full color represent genes with *p*_adj_ below 0.05 (dashed grey horizontal line, -log_10_ *p*_adj_ = 1.3) and a FC above 1.41 (vertical dashed grey lines, log_2_ FC = ± 0.5). Genes with non-significant variation are displayed with reduced opacity.

### CIRBP mRNA targets shift from housekeeping- to stress response-related functions upon infection

As CIRBP acts as a regulator of gene expression by binding its target mRNAs, we compared the sets of CIRBP targets between non-infected and infected cells by RNA-immunoprecipitation of CIRBP followed by Illumina sequencing of co-precipitated mRNAs (RIP-seq) (Table S4). From a total of 242 identified mRNA targets, a common core of 146 genes were similarly enriched in CIRPB immunoprecipitations (IPs) in both conditions (Figure 5B). By contrast, two distinct sets of mRNAs were preferentially bound by CIRBP depending on infection; 28 targets were preferentially enriched in non-infected cells (Figure 5B, blue dots; Figure 5C), and 68 others were preferentially enriched in infected cells (Figure 5B, pink dots; Figure 5D), likely reflecting a reallocation of CIRBP regulatory activity on cellular mRNAs in response to infection. Comparison of the sets of CIRPB-bound targets by gene ontology overrepresentation analysis highlighted an overall shift in the functions of their products between the two conditions (Figure S8). Enrichment for housekeeping functions (transcription, organization of the cytoskeleton (especially centrosomes) or metal cation binding was reduced upon infection, in favor of functions associated with cellular response to stress, DNA damage response and repair, ubiquitination and proteolysis, and regulation of metabolic processes. The observed changes in the set of CIRBP targets suggest that infection-induced control of CIRBP activity contributes to the coordinated regulation orchestrating the cellular response to infection-induced stress.

To investigate a potential role of CIRBP in the regulation of its targets during infection, we have intersected the set of 242 CIRBP targets from RIP-seq analysis with data of differential expression between non-infected and 10-h-infected cells from RNA-seq (Figure 5E, Table S4E), Ribo-seq (Figure S9A, Table S4E), translational efficiency (TE, Figure 5F, Table S4E), or differential LSVs (Figure S9B, Table S4F). CIRBP targets were not enriched for a differential use of LSVs (Fisher’s test *p* = 0.2), suggesting that CIRBP itself does not behave as a regulator of AS in this context (Figure S9B). By contrast, there was a strong enrichment of CIRBP mRNA targets for differential RNA expression (Fisher’s test *p* = 2.7 × 10^-9^), and most strikingly for increased mRNA abundance upon infection (Figure 5E). Conversely, there was an enrichment for transcripts with reduced TE in infected cells among CIRBP targets (Figure 5F) (Fisher’s test *p* = 1.4 × 10^-6^). These two observations were mirrored in the ribosome footprint plot (Figure S9A) (Fisher’s test *p* = 2.4 × 10^-6^): genes that are translated proportionally to their increased mRNA abundance display increase footprint levels, whereas those that are translationally repressed while not varying in mRNA abundance display reduced footprints (Figure S9A). These data suggest that the modulation of CIRBP abundance, subcellular localisation and/or function during infection impacts the stability or translation of its targets in a proportion that contributes to the host gene expression response to infection.

## Discussion

A broad range of damaging challenges trigger the activation of coordinated cell stress response pathways that integrate signaling cascades and gene expression regulation. Among central players in the coordination of cell responses, CIRBP has previously been documented to regulate the expression of mRNAs involved in the modulation of the cell cycle and of inflammation, following exposure to multiple types of abiotic stressing agents (heat stress, UV or ionizing radiations, hypoxia, oxidative or osmotic stress, nutrient deprivation) or to viral infections (3, 4, 12). Our findings extend the role of CIRBP to the control of cell autonomous responses to biotic stress caused by bacterial PFTs upon a bacterial infection, with downstream effects on the modulation of infection efficiency in a cell monolayer (Figure 6). In spite of the distinct nature of the initial stressors, our results highlight that the mechanisms of regulation of CIRBP by AS are shared between the previously-described response to heat stress (7, 9) and to PFT-dependent membrane damage. The exact nature of the unbalance that is sensed upon membrane permeation is currently unclear, as discriminating effects of intra/extracellular cation concentrations from that of a perturbation of the plasma membrane potential constitutes a technical hurdle.

**Figure 6.**
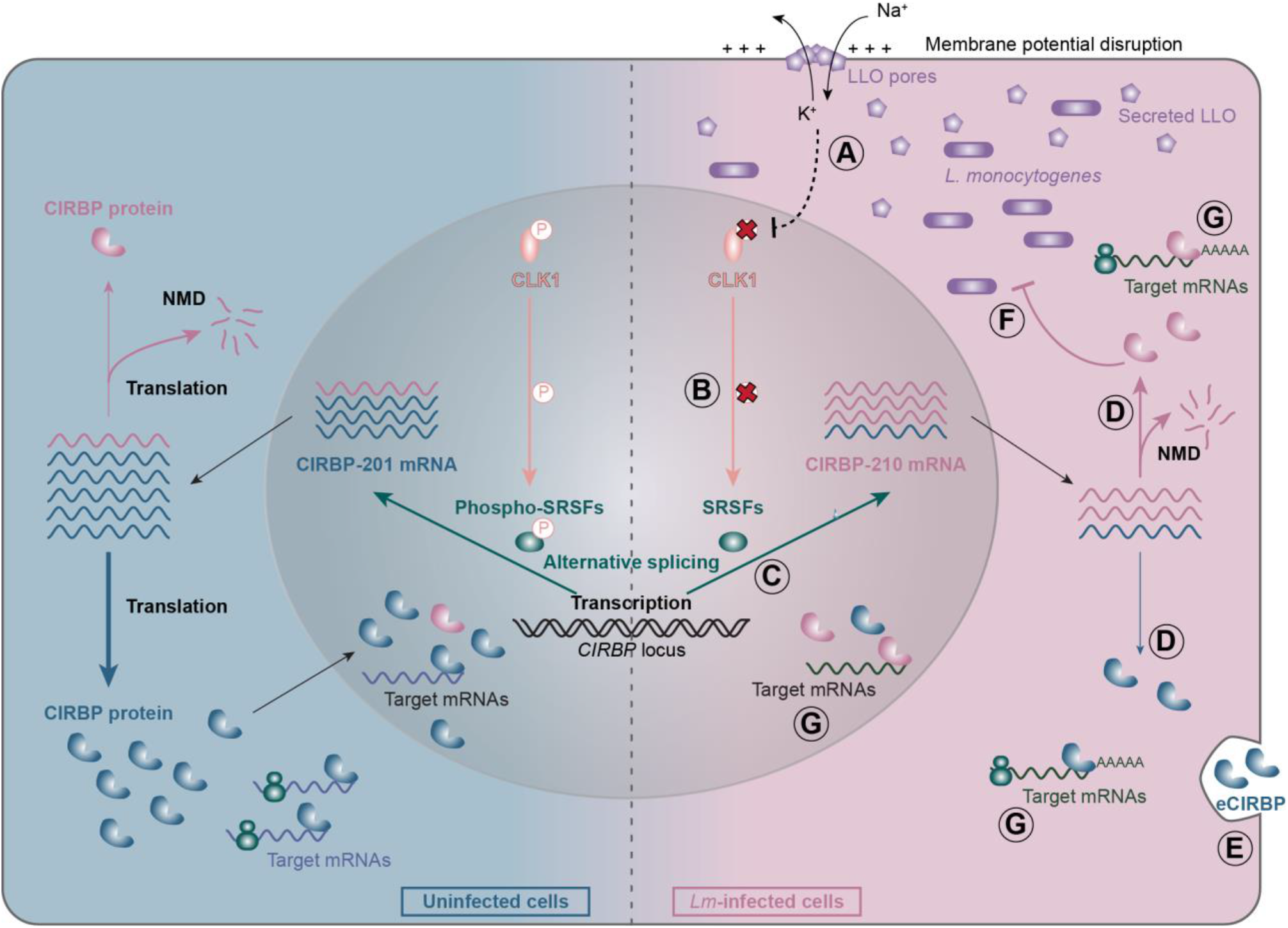
Model of CIRBP regulation in response to *Listeria monocytogenes* infection. In uninfected cells (left, blue background), the splicing of CIRBP pre-mRNA is in favor of the CIRBP-201 isoform, which is translated into the canonical 18-kDa CIRPB protein that regulates the expression of a subset of human mRNAs. In *Lm*-infected cells (right, pink background), upon LLO-induced membrane damage (A), CLK1 activity is inhibited, which results in a decreased phosphorylation of SR proteins (SRSFs) (B). This causes a shift in CIRBP splicing in favor of the CIRBP-210 isoform (C). As this isoform is a target of NMD, it is unstable, which results in reduced overall mRNA abundance and translation (D). Secretion of CIRBP (eCIRBP) in response to LLO-induced cell damage also likely contributes to reduce the pool of preexisting CIRBP protein (E). Expression of CIRBP-210 displays anti-bacterial effects, contrary to that of CIRBP-201 (F). The subset of CIRBP-bound mRNAs in infected cells also differs from uninfected cells (G).

Following stress sensing, the CLK1 kinase appears to play a key role as an integrator of distinct types of damaging challenges by controlling the acvtivity of SR splicing factors, although the signalling pathways bridging stress sensing with the control of CLK1 activity are still elusive. CLK1-dependent inhibition of SR proteins reorients the splicing of specific cellular tanscripts, including CIRBP and CLK1 itself, towards the production of alternatively processed mRNAs (7, 44). Pre-mRNA processing of CIRBP is then shifted from the use of a canonical 3’-UTR to an alternative 3’-UTR that contains a spliced intron. Upon splicing, deposition in the 3’-UTR of an exon junction complex that cannot be removed by translating ribosomes render this type of transcripts sensitive to NMD, and thus unstable (46). The stress-induced AS of CIRBP transcripts in favor of an NMD-sensitive RNA isoform thus lowers CIRBP mRNA expression, including in response to LLO-dependent stress during *Lm* infection and more generally to the stress induced by several classes of bacterial pore-forming toxins. CIRBP protein levels are also down-regulated by LLO-induced stress. The release of eCIRBP following LLO-dependent membrane damage likely contributes to this reduction, although we cannot rule out that the stability of the protein is also impacted by LLO activity. Following eCIRBP release, the reduced levels of CIRBP mRNAs in infected cells would oppose to the replenishment of CIRBP protein stores.

Besides the control of CIRBP AS by the CLK1-dependent activity of SR splicing factors, it has been previously reported that modulation of m^6^A modification in the 6^th^ exon of CIRBP influenced its AS, with a positive effect of methylation on the expression of the long protein-coding, CIRBP-225 isoform (12). In the present study, CIRBP-225 was expressed at very low level in the LoVo cell line and not significantly affected by infection. However, given the involvement of the splice junction in exon 6 in both cases, it is to be wondered whether CIRBP-201 to -210 AS might also depend on a differential methylation in exon 6 for the selection of the acceptor sites in exons 7a and 7b, respectively. Deeper investigation would also be required to determine if m^6^A modification of CIRBP mRNA and SR protein activity are independent, additive mechanisms resulting in distinct forms of CIRBP AS in different contexts, or if one of them would condition the other.

In addition to quantitative effects on CIRBP mRNA stability due to NMD, the isoform switch in CIRBP that is observed upon *Lm* infection also impacts the sequence of the encoded protein products. It is thus expected that CIRBP-210 product gradually replaces CIRBP-201 product after eCIRBP release. Discriminating the impact of protein sequence change from quantitative effects has not been addressed in the present study, nor in the existing literature to the best of our knowledge, and is worthy of detailed investigation in the future. The divergence between CIRBP-201- and -210-encoded proteins only affects the last four residues in the sequence, but entails the loss of a putative phosphorylation site previously hypothesized to participate in the nucleocytoplasmic partitioning if CIRBP (35), which is functionally relevant since the stress-induced translocation of CIRBP to the cytoplasm impacts it ability to bind and regulate its target genes (3). Hence, a change in the localization or functionality of the CIRBP-210 product could take a share in the differential effect of the two isoforms on the binding to and regulation of CIRBP target mRNAs and on infection, on top of purely quantitative effect due to altered RNA stability.

Our observation that expression of CIRBP-201 is rather pro-bacterial is consistent with a previous report where both the dominant CIRBP-201 and the low abundance -225-like protein-coding isoforms were found to have proviral effects, independently of their respective localizations (predominantly nuclear for the former, cytoplasmic for the latter) (12). Both findings hint at a detrimental effect of expression of canonical CIRBP on the ability of epithelial cells to face infectious challenges. However, isoform-based regulation differs between the two studies. During viral infections, quantities of short-size CIRBP-201 (or -210, although it was not addressed in that study) products (CIRPB-S) remained stable, while levels of long-sized CIRBP-225 products (CIRBP-L) were reduced due to the above-mentioned loss of m^6^A modification in the 6^th^ exon. By contrast, upon *Lm* infection we observed a reduction of CIRBP-201 levels and of CIRBP-S protein levels, while expression of CIRBP-225 remained invariably low (Table S3G). These differences in expression may reflect intrinsic differences between the two model cell lines in the set of isoforms they express, or distinct mechanisms of regulation of CIRBP upon cellular stresses on the one hand, and viral infection on the other hand. Comparatively addressing the differential contribution of CIRBP-201, -210 and - 225 isoforms to the control of viral and bacterial infections, respectively, would deserve closer attention in the future.

In response to *Lm* infection, CIRBP overall activity is impacted not only by the regulation of its abundance and the nature of its isoforms, but also likely by its localization (cytoplasmic, nuclear, or in membraneless organelles) and post-translational modifications, as previously reported in response to other stressors (5). The two latter levels of regulation will deserve close investigation, as they can strongly condition the ability of CIRBP to bind and regulate its target genes during infection. Although a common core of CIRBP targets identified in RIP-seq was similarly bound in infected and uninfected conditions, preferential binding of CIRBP to a subset of mRNAs in infected cells likely reflects a modulation of its affinity for its binding sites or of its localization. The loss of a putative phosphorylation site in the product of CIRBP-210 could for instance have an impact on CIRBP binding features, or on its ability to reach its targets due to an altered subcellular distribution. Alternatively, LLO-induced stress could affect CIRBP phosphorylation or methylation independently of its protein isoform (3, 35).

During infection, a consistent fraction of CIRBP targets tend to display increased mRNA levels and, as a consequence, be more translated. This could proceed from a positive impact of CIRBP on the stability of its target mRNA during infection or on the contrary, for targets that are shared between the two conditions, from a repressive role of CIRBP in uninfected cells that would be alleviated upon infection due to reduced CIRBP abundance. Among the mRNAs that are preferentially bound by CIRBP in infected cells, BAG6, LTN1, WDR26 and ZEB1 are upregulated. Interestingly, the proteins encoded by these genes have functions related to the control of cells death (BAG6, WDR26), ubiquitin-dependent proteolytic degradation of proteins in response to stresses (LTN1, WDR26) or regulation of immunity (ZEB1, BAG6), exemplifying a larger pool of CIRBP infection-specific targets with a function related to cellular response to stress. Given their cellular roles, upregulation of these factors could be part of a CIRBP-mediated cellular response to better accommodate LLO-induced stress and respond to infection.

Another set of CIRBP targets —that are bound both in infected and non-infected cells— is translationally repressed during infection. This repression might reflect a relocation of CIRBP and of its mRNA targets to translationally-inactive stress granules, as previously shown upon osmotic or heat shock (3). Of note, several of the translationally-repressed targets of CIRBP belong to the translational regulon (EEF2, EIF4B, PABPC1, RPL3), which we had previously reported to be translationally-repressed upon LLO-induced infection stress as part of a response contributing to dampen bacterial loads (22). A typical feature associated with their co-repression was the presence of a 5’-terminal oligopyrimidine motif in their 5’-UTR. Whether CIRBP binding to these targets is coincidental, or whether it is actively contributing to their addressing to stress granules and translational repression would deserve to be untangled in future studies.

Beyond human cells, roles for CIRBP homologs have been previously described in the adaptation of a broad range of multicellular eukaryotes to various types of environmental changes. In reptiles, CIRBP has been reported to be involved in temperature-dependent sex-determination, in line with its role during moderate cold- or heat-exposure in mammals (7, 47). In plants, CIRBP homologs of the glycine-rich RNA-binding protein (GRP) family are involved in the control of innate immune responses to bacterial invaders as well as in cold adaptation and resistance to high salinity or drought (48–50). These similarities of contribution to the coordination of responses to biotic and abiotic stresses across distant kingdoms of the Eukaryota hints at an ancient, conserved function of this family of proteins as key players in the coordination of defenses against pathogens and damaging environmental challenges.

## Supplementary data

Supplementary data are available as Supplementary Figures S1 to S9 and Supplementary Tables S1 to S4.

## Data availability

All novel sequencing data discussed in this publication have been deposited in the ArrayExpress database (https://www.ebi.ac.uk/biostudies/arrayexpress) and are accessible under accession numbers <u>GSE225417</u> for long-read sequencing, <u>GSE225516</u> for RIP-seq. Previously published transcriptomic data obtained with Illumina sequencing are available from the European Nucleotide Archive under accession number <u>PRJEB26593</u> (22). All other data is contained within the manuscript and/or supplementary files.

## Funding

Research at IBENS received support under the program “Investissements d’Avenir” managed by the French national research agency (ANR-10-LABX-54 MemoLife and ANR-10-IDEX-0001-02 PSL). Research in AL’s group was supported by Mairie de Paris (programme Émergence(s) Recherche médicale). The GenomiqueENS core facility was supported by the France Génomique national infrastructure, funded as part of the “Investissements d’Avenir” program managed by the French national research agency (ANR-10-INBS-09). MC received doctoral fellowships from the MESRI (Ministère de l’enseignement supérieur, de la recherche et de l’innovation) and from LabEx MemoLife.

The authors declare no competing interests. The funders took no part in the orientation of research.

## Acknowledgements

We are grateful to Hervé Le Hir, Lucia Morillo, Nadia Ruiz-Gutierrez, Isabelle Barbosa, Laurent Jourdren, Sophie Lemoine, Dominique Weil, Lionel Navarro, Laetitia Vincensini, Brice Sperandio, Caroline Peron-Cane, Thomas Petit and Simon Guette-Marquet for precious scientific discussion, methodological advice and kind help, and to Olivier Disson for reagents sharing.

## Author contributions

AL initiated and supervised the study. MC and AL devised the research hypotheses and methodology, and interpreted results. MC performed most experiments with occasional assistance by AK and AC, who she supervised. VBe provided RNA samples for long-read RNA-seq and methodological advice. AM and CSB prepared sequencing libraries and performed sequencing at the École normale supérieure genomics core facility. MC analysed sequencing data with advice and input from VBoe and MTC. MC and AL wrote the manuscript.

## Supplementary material

**Figure S1.**
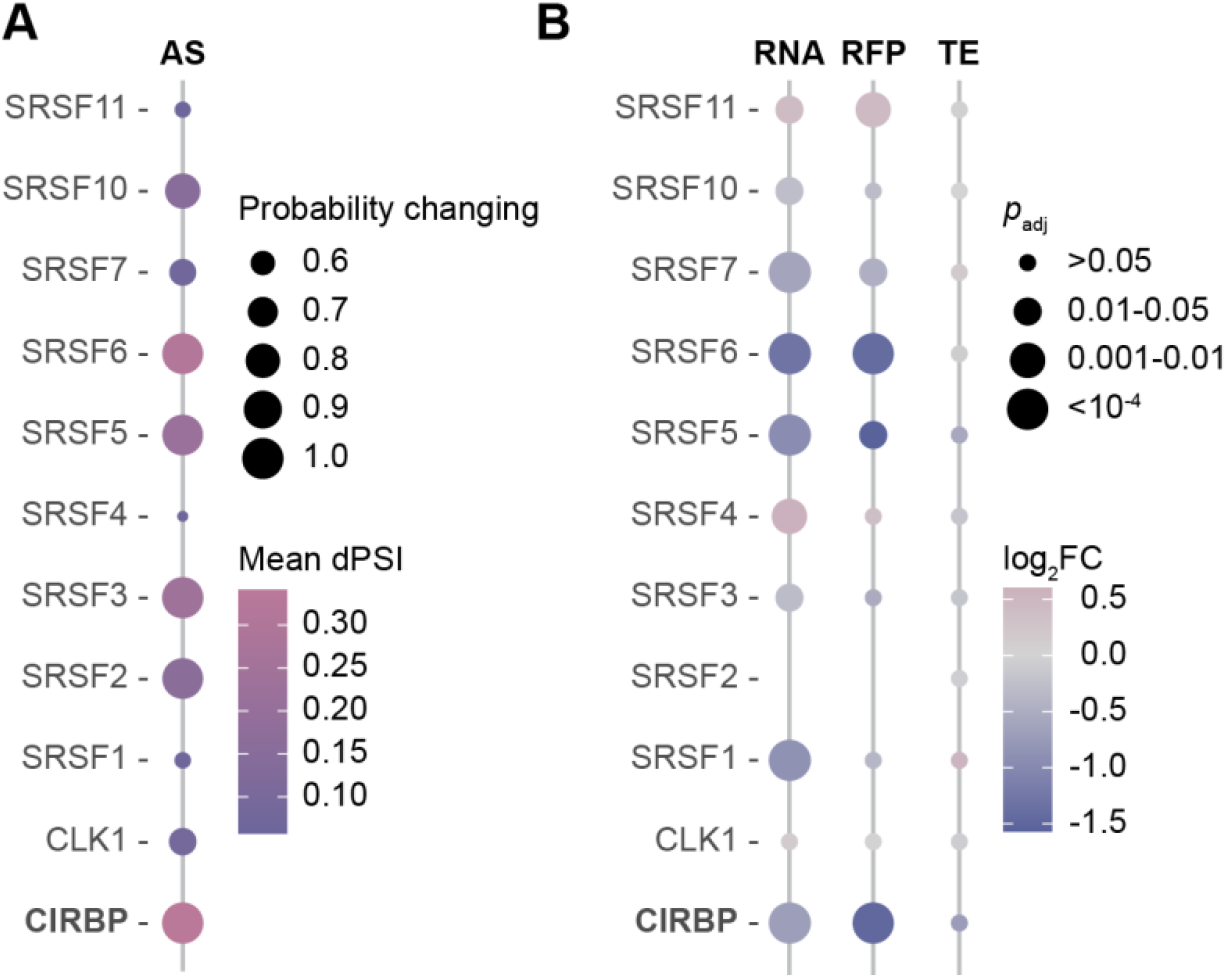
Expression of human genes affected by isoform-based regulations upon 10-h infection with *Listeria monocytogenes*. (A-B) Dot plot highlighting (A) alternative splicing events (dPSI and probability of changing) for a subset of genes of interest at 10 hpi, and (B) their corresponding genes expression in terms of differential mRNA abundance (RNA), ribosomal occupancy (ribosome footprints, RFP) and translation efficiency (TE) when compared to uninfected cells. log_2_FC (fold change); *p*_adj_, adjusted *p*-value [DESeq false discovery rate (FDR)]. Data from three independent replicates of RNA-seq and Ribo-seq short-read cDNA sequencing at 0, 2 and 5 hpi; 2 replicates at 10 hpi.

**Figure S2.**
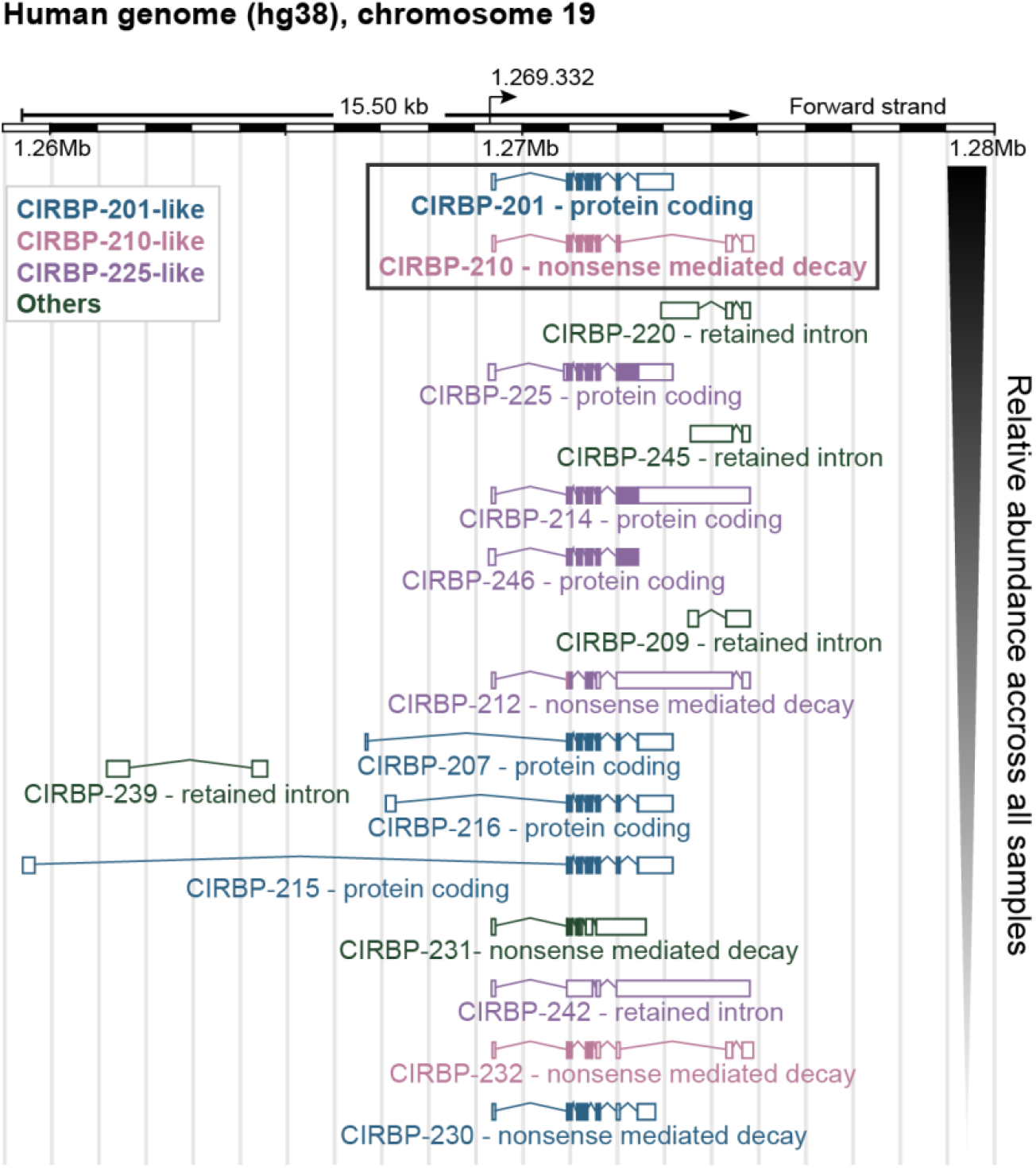
Map of CIRBP isoforms detected in LoVo cells by long-read sequencing. Assigned isoforms that represented more than 0.5% of overall transcripts in long-read sequenced samples after Bambu-ISAR analysis are represented by decreasing abundance. CIRBP splice variants are grouped into four categories based on LSVs involving the junction of position 1, 272, 051 in exon 6 with downstream sequences. (*a*) In CIRBP-201-like isoforms, in blue, the acceptor site is located at position 1, 272, 427 (exon 7a). The prototype version, CIRBP-201, encodes a 172-amino-acid protein. (*b*) In CIRBP-210-like isoforms, in pink, the acceptor splice-site is shifted to position 1, 274, 307 (exon 7b). The prototype CIRBP-210 harbors a spliced exon between exons 7b and 8 in its 3’-UTR, which make it an NMD target and thus unstable. The protein it encodes only has a few amino acid differences compared with CIRBP-201. (*c*) In CIRBP-225-like transcripts, in purple, no splicing occurs at position 1, 272, 051, leading to an intron retention. Most CIRBP-225-like isoforms encode elongated versions of the CIRBP protein (292 amino acids for CIRBP-225). (*d*) Other isoforms. The position of the common transcription start site detected for complete nanopore reads aligning with CIRBP-201- and CIRBP-210-like transcripts in our datasets is indicated with a black arrow.

**Figure S3.**
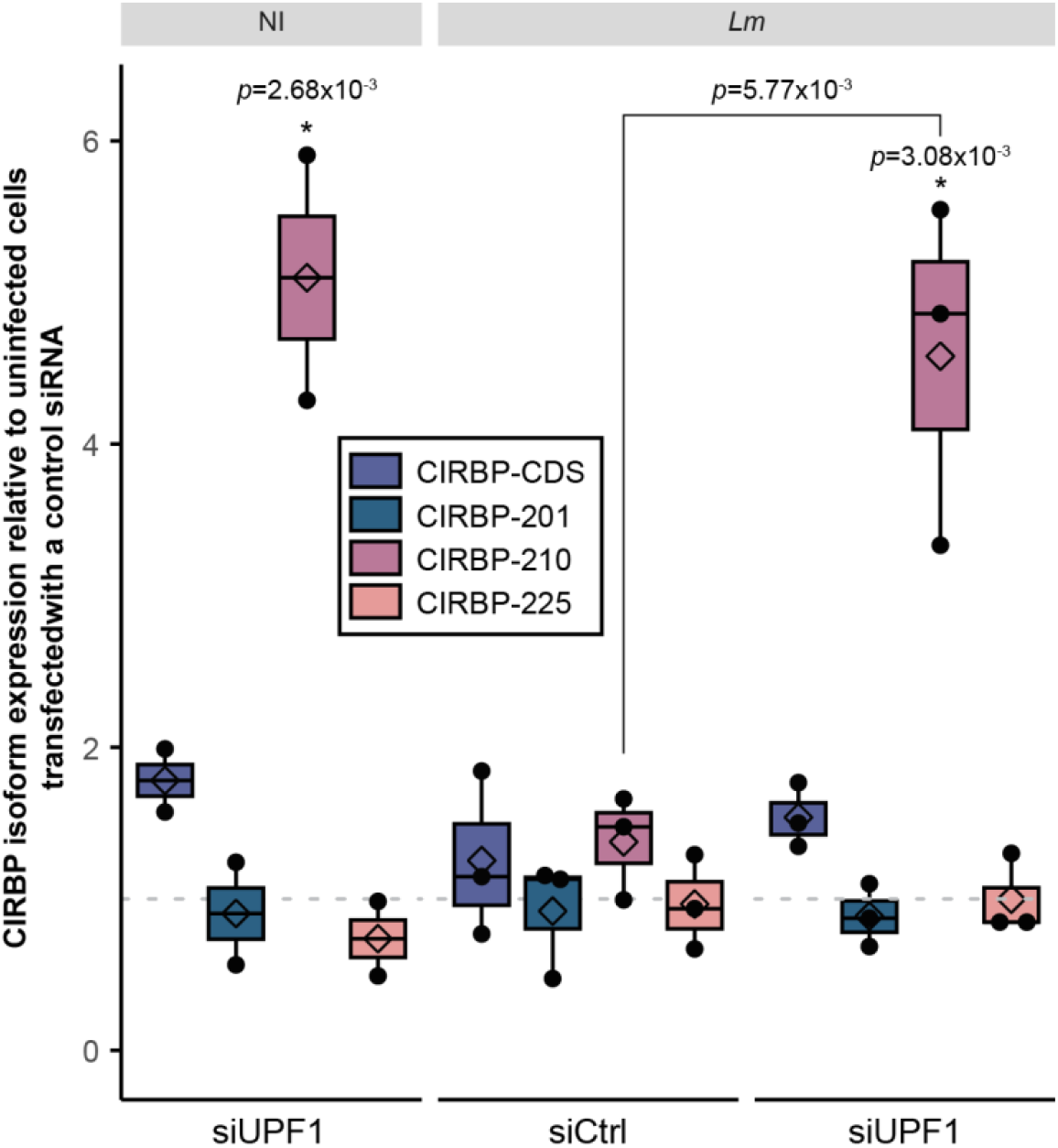
Expression of CIRBP isoforms in conditions of repression of the NMD and/or of infection by *Listeria monocytogenes*. Total CIRBP mRNAs (CIRBP-CDS) or each one of the isoforms CIRBP-201, -210 or -225 were quantified by RT-qPCR in LoVo cells transfected with a siRNA targeting UPF1 (siUPF1) or a scrambled siRNA (siCtrl), either non-infected (NI) or in cells infected for 10 h with *Lm*, using isoform-specific primers, and normalized to GAPDH. Expression levels are expressed relative to the level of each transcript in uninfected cells transfected with the scrambled siRNA. Boxplots represent medians and quartiles of data from four independent experiments. One way ANOVA followed by post-hoc Tukey’s test was used for statistical testing between conditions.

**Figure S4.**
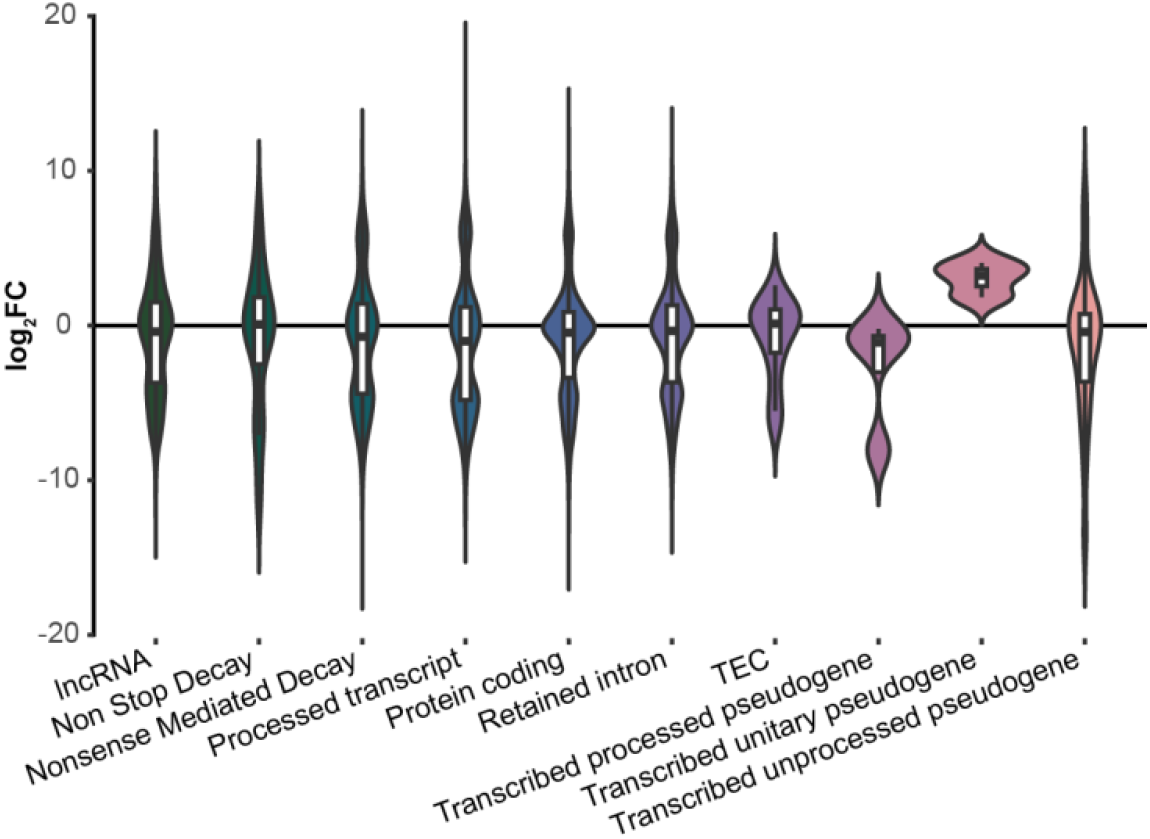
Distribution of isoform switches according to isoform biotypes. Violin plot of the log_2_ fold change (FC) distribution of the switching isoforms for each isoform biotype represented in BioMart: long non-coding RNA (lncRNA), non-stop decay targets, nonsense mediated decay targets, stable protein coding isoforms, isoforms with a retained intron, so-called “to be experimentally confirmed” (TEC) isoforms, transcribed processed pseudogenes, transcribed unitary pseudogenes and transcribed unprocessed pseudogenes.

**Figure S5.**
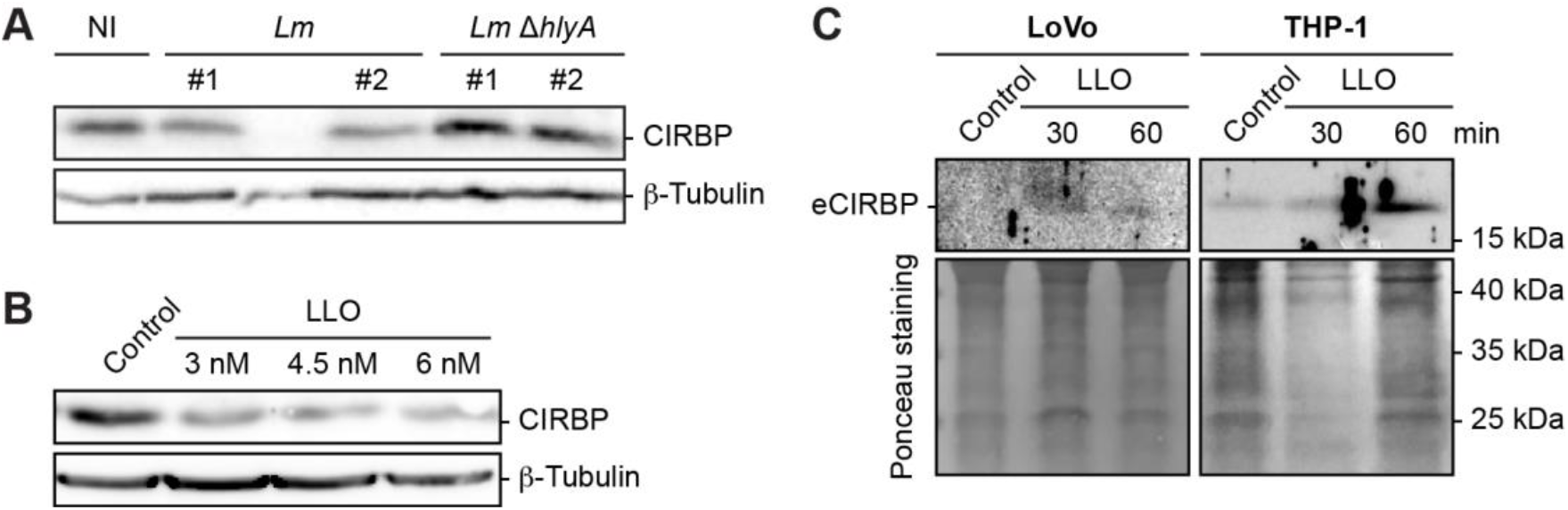
CIRBP protein levels are reduced in cells exposed to LLO. (A-B) Detection of CIRBP by immunoblot in LoVo cells infected for 10 h with WT *Lm* or *Lm* Δ*hlyA*, compared to uninfected cells (NI) (A), or in cells treated for 1 h with increasing concentrations of LLO, compared to untreated cells (B). (C) Detection of extracellular CIRBP (eCIRBP) in the supernatants of LoVo or THP-1 cells exposed to LLO. Note that the first lane in (A) and the fifth lane in (C) are underloaded, which however does not affect conclusions.

**Figure S6.**
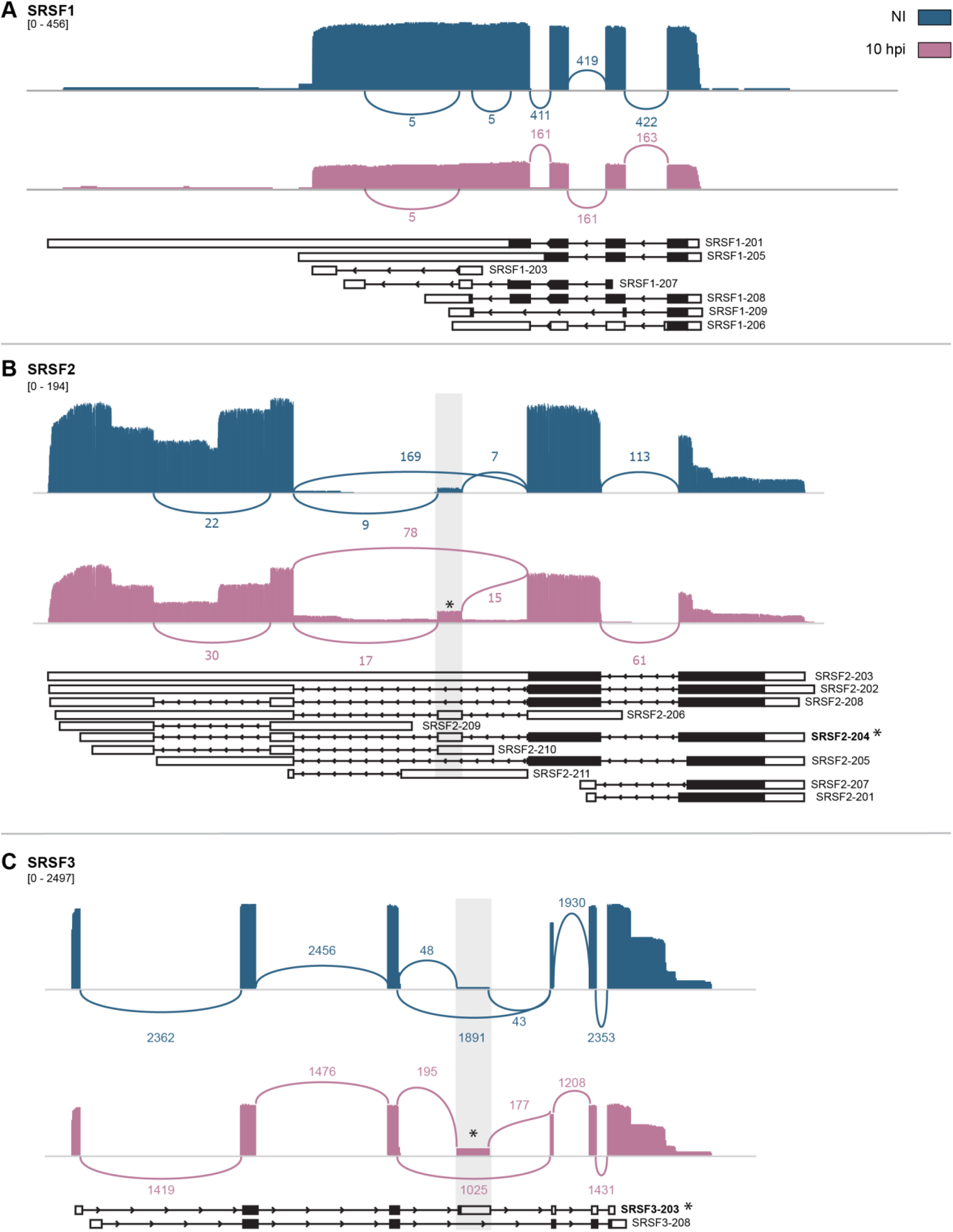
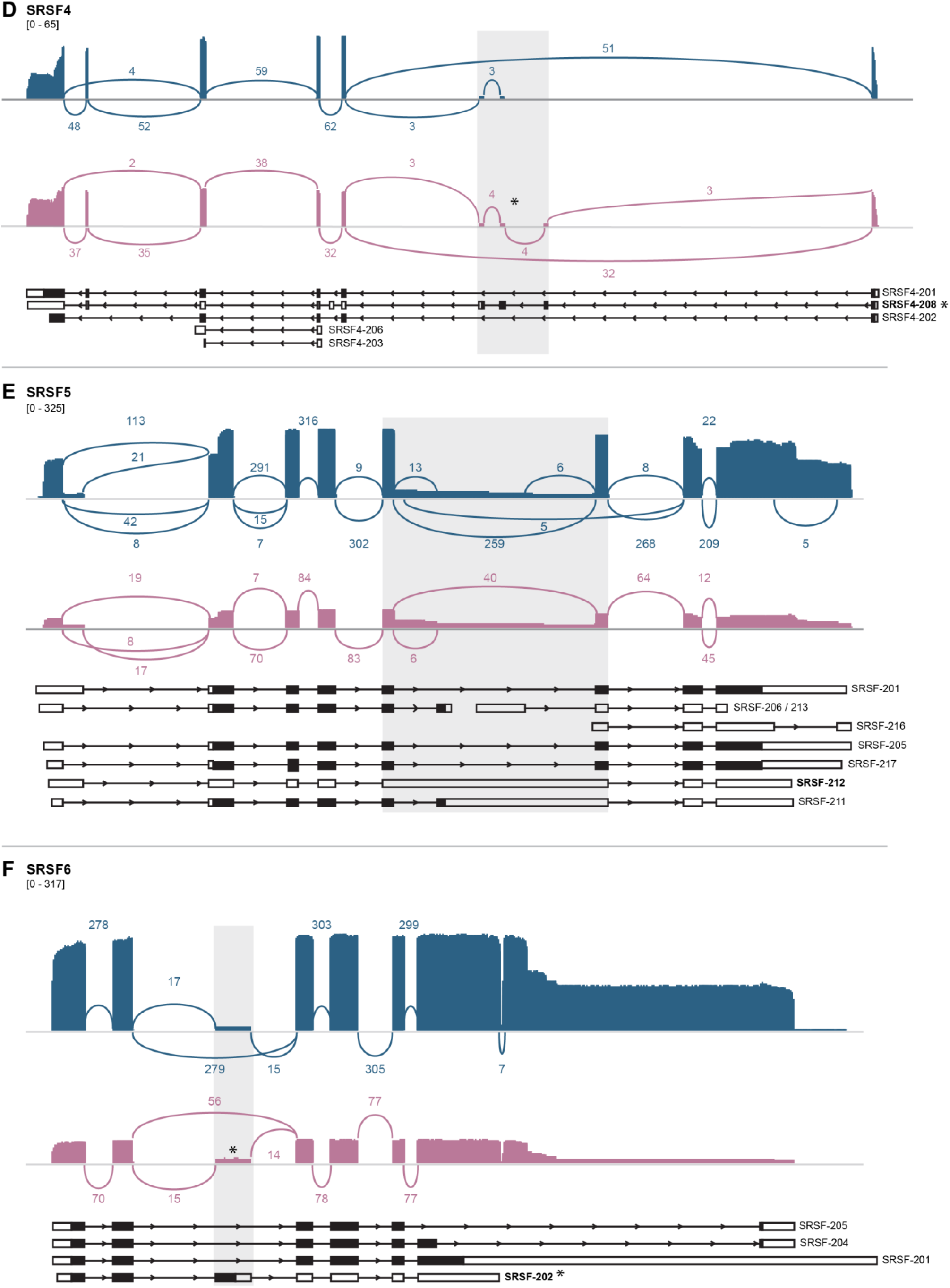
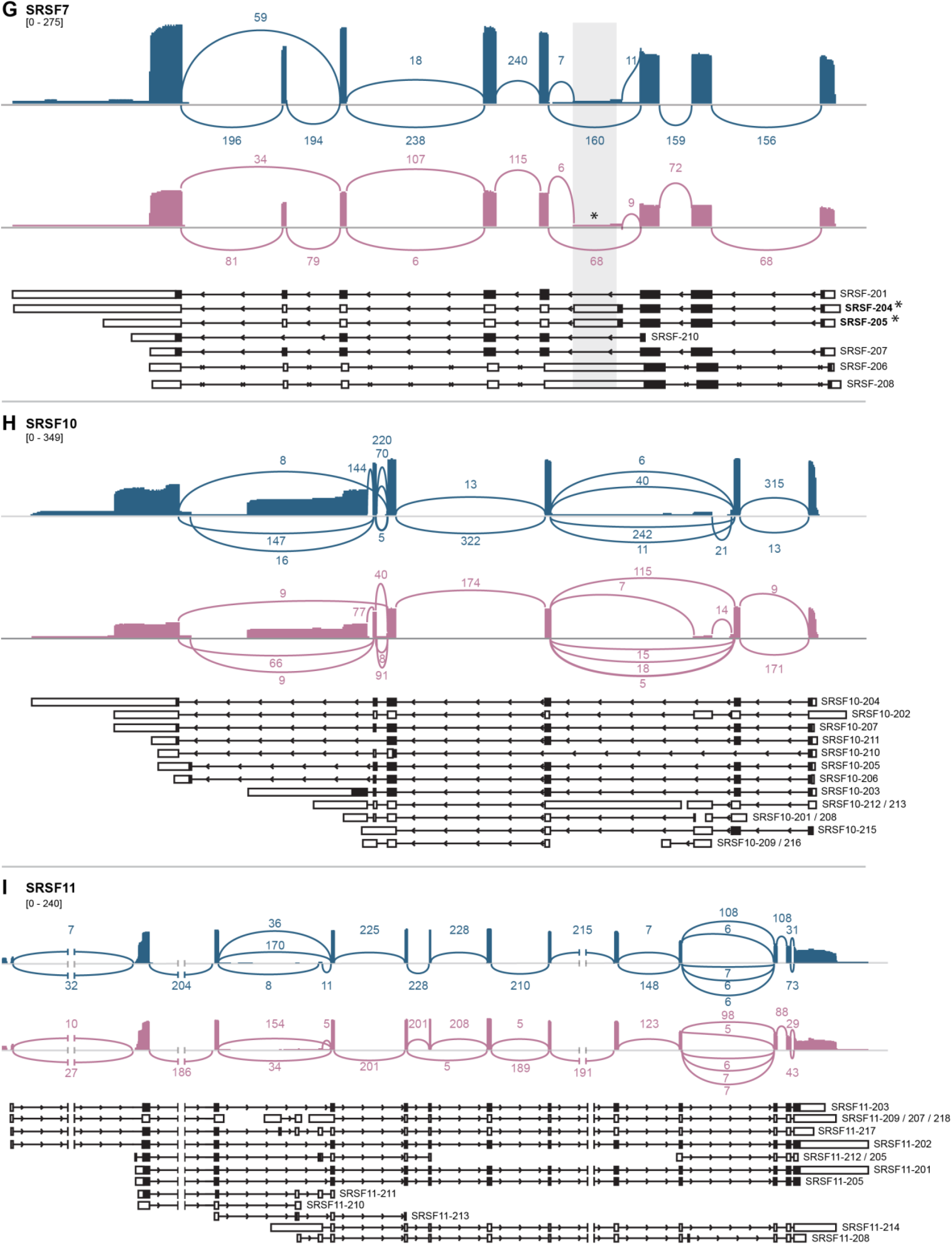
Alternative splicing events affect the expression of a subset serine-arginine rich splicing factors (SRSFs) upon *Listeria* infection. (A-I) Sashimi plots of a subset of SRSF genes, representing exon coverage and exon junctions in uninfected (blue) and *Lm*-infected cells at 10 hpi (pink). Average values of read counts per genomic position normalized for library size, and exon-junction coverages for each junction, are represented (3 independent replicates at 0, 2, 5 hpi; 2 at 10 hpi). The range for minimum and maximum normalized read counts used as a scale for each SRSF is indicated between brackets, and allows the visualization of overall expression changes between the two conditions. Transcript isoforms identified by mapping long reads to Ensembl annotated transcripts are represented for each gene, with coding sequences displayed as black boxes, and untranslated regions as open boxes. Regions affected by alternative splicing events are highlighted in light grey. Stars indicate the most representative alternative splicing event that led to the inclusion of poison exons due to the presence of premature termination codons (SRSF2, SRSF3, SRSF4, SRSF6 and SRSF7). Other such events occurred, in SRSF1 for instance, but were less obviously visualized due to a multiplicity of coexpressed isoforms and to the low stability of transcripts containing poison exons. Note that precise isoform attribution is sometimes hampered due to non-exhaustivity of current transcript annotations (see for instance SRSF3-208, the 5’-end of which does not correspond to the transcript that is actually produced in LoVo cells); no corresponding transcript with a matching set of exons is currently available in Ensembl).

**Figure S7.**
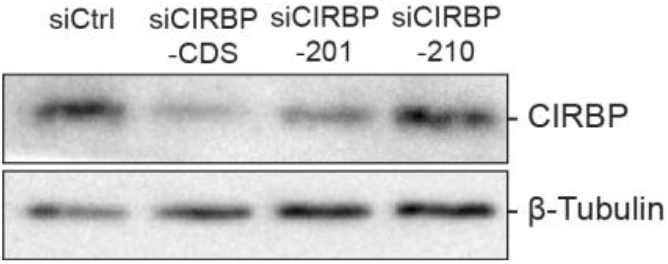
CIRBP protein expression upon CIRBP knock-down. Assessment of CIRBP protein expression by immunoblotting in uninfected cells transfected by siRNA targeting the common CDS between CIRBP-201 and CIRBP-210 (siCIRBP-CDS), the 3’-UTR of CIRBP-201 specifically (siCIRBP-201) or the 3’UTR of CIRBP-210 specifically (siCIRBP-210). Compared to cells transfected with a scrambled siRNA (siCtrl), a decrease in CIRBP protein levels was observed upon treatment with siCIRBP-CDS and siCIRBP-201, but not with siCIRBP-210, consistently with CIRBP-210 not being expressed in uninfected cells.

**Figure S8.**
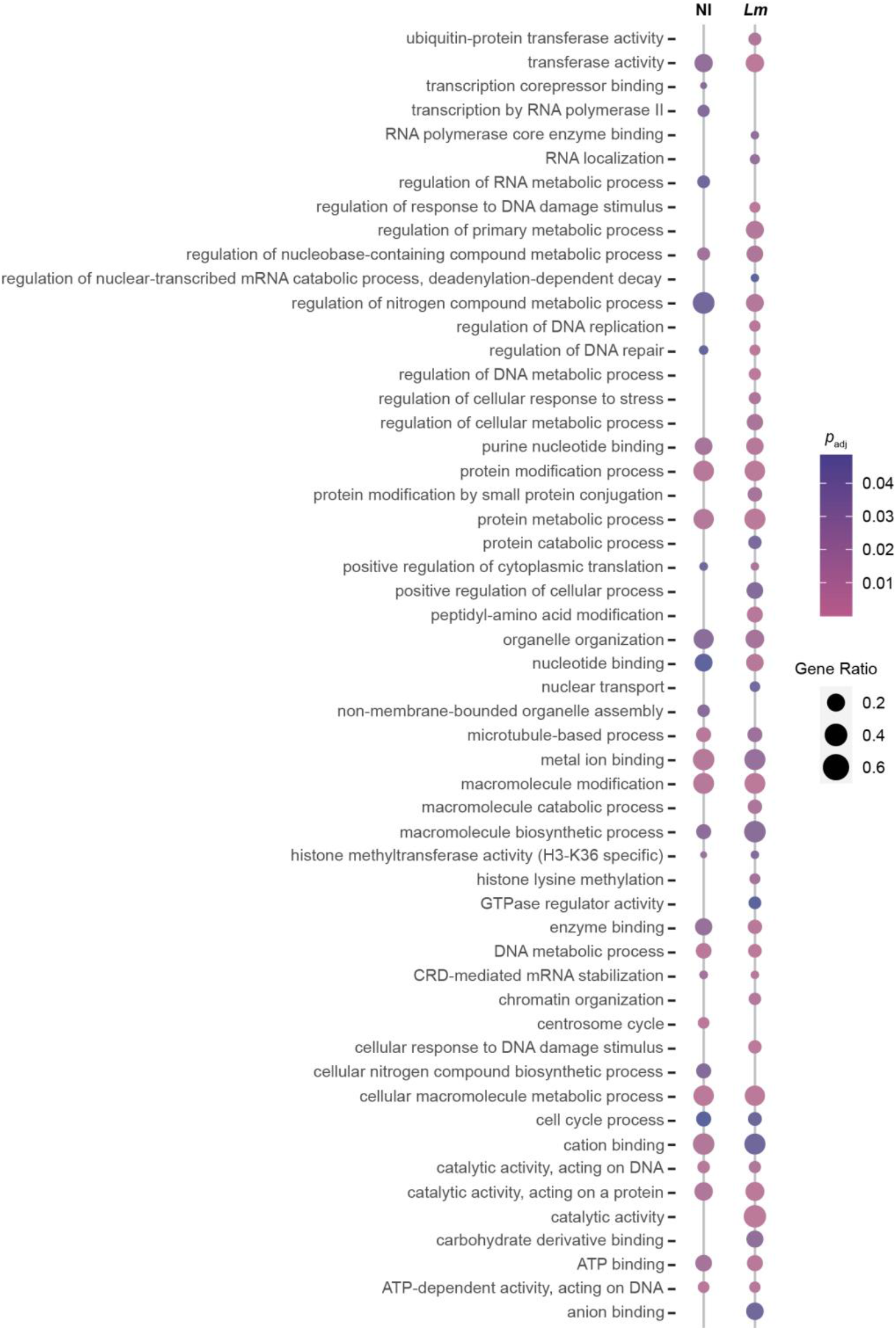
Main biological processes in which products of CIRBP targets are involved. Over-representation analysis of GO Biological Process terms for CIRBP targets identified by RIP-seq analysis, either in non-infected (NI) LoVo cells, or in cells infected by *Lm* for 10 h. Only transcripts enriched in the CIRBP RIP when compared by differential expression analysis either to the input, or to control IgGs were kept as targets (223 genes in total). *p*_adj_, adjusted *p*-value.

**Figure S9.**
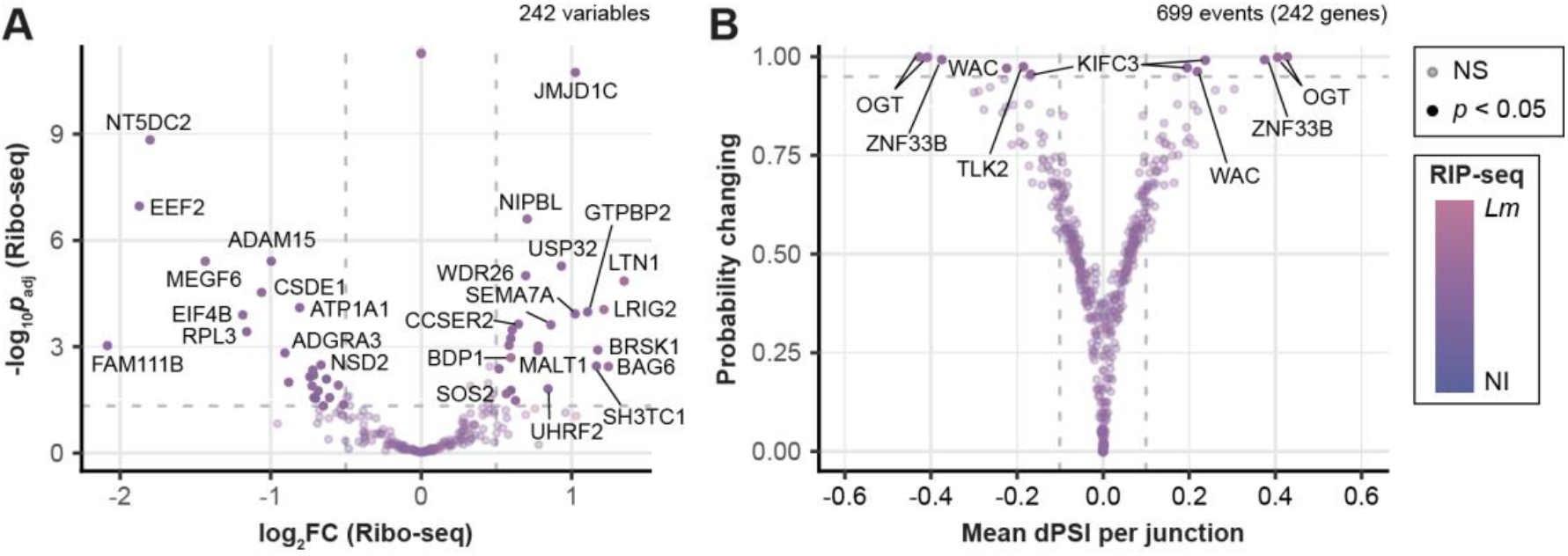
Variations in translation and in alternative splicing among CIRBP mRNA targets. Volcano plots highlighting changes in mRNA levels (Ribo-seq data; A) and LSVs (output data from MAJIQ; B) for the 242 CIRBP targets, in cells infected with *Lm* for 10 h compared to non-infected cells. The color of data points ranging from blue to pink reflects the Δlog_2_FC of RIP enrichment in infected (*Lm*) versus non-infected (NI) cells. Data points colored in full color represent genes with *p*_adj_ below 0.05 (dashed grey horizontal line, -log_10_ *p*_adj_ = 1.3) and a FC above 1.41 (vertical dashed grey lines, log_2_ FC = ± 0.5). Genes with non-significant variation are displayed with reduced opacity.

**Table S1.** Bacterial strains used in this study.

| Strain genotype | Culture medium | MOI | Reference |
| --- | --- | --- | --- |
| *L. monocytogenes* LL195 | BHI | 30 | Weinmaier et al., 2013 |
| *L. innocua* [pGM30-*inlA*(LRRs-IR-SPA)] | BHI + Cm 7µg/mL | 100 | Lecuit et al., 1997 |
| *L. monocytogenes* LL195 Δ*hlyA* | BHI | 30 | Besic et al., 2020 |

**(B) Bacterial strains used for the purification of PFTs**
| Strain genotype | PFT | Culture medium | Reference |
| --- | --- | --- | --- |
| *E. coli* Top10(DE3) [pET29b-LLO-6His] | LLO | LB + Km (30 µg/ml) | Glomski et al., 2002 |
| *E. coli* Top10 [pET29b-LLO<sub>C484A</sub>-6His] | LLO<sub>C484A</sub> | LB + Km (30 µg/ml) | Ribet et al., 2010 |
| *E. coli* Top10 [pET29b-LLO<sub>W492A</sub>-6His] | LLO<sub>W492A</sub> | LB + Km (30 µg/ml) | Ribet et al., 2010 |
| *E. coli* Top10 [pET29b-LLO<sub>Y206A</sub>-6His] | LLO<sub>Y206A</sub> | LB + Km (30 µg/ml) | Ribet et al., 2010 |
| *E. coli* DH5a [pTrcHisA-PFO] | PFO | LB + Ap (100 µg/mL) | Farrand et al., 2010 |
| *E. coli* DH5a [pTrcHisA-SLO] | SLO | LB + Ap (100 µg/mL) | Farrand et al., 2010 |
| *E. coli* NEB5a [pET22b-PA-6His] | ALY | LB + Ap (100 µg/mL) | Iacovache et al., 2006 |

Columns indicate the identifier of the strain, its species background and plasmid, the produced PFT, the medium used for culture and the source reference. LB, Luria-Bertani medium; Km, kanamycin; Ap, ampicillin.

**Table S2.**
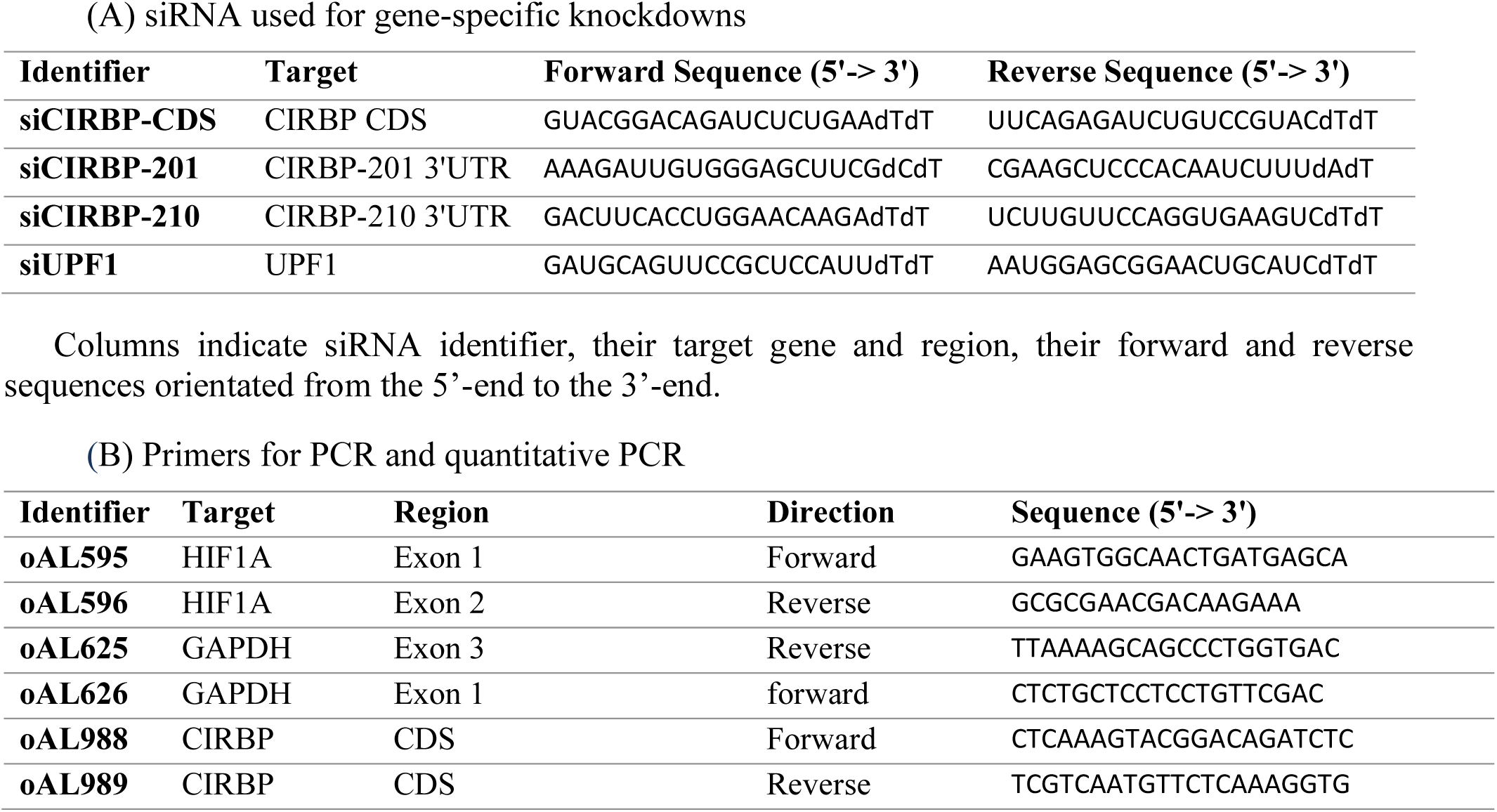
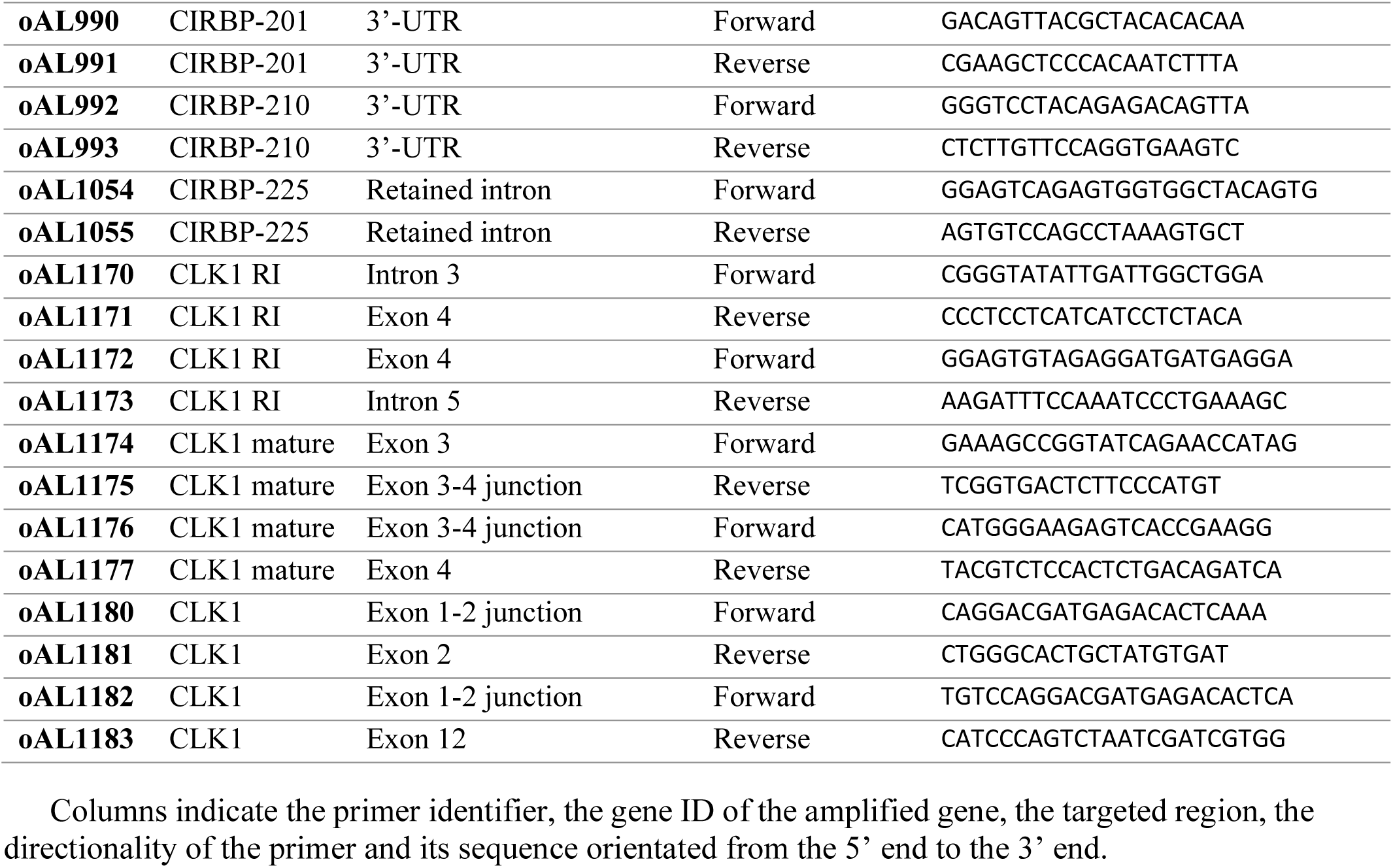
Oligonucleotides used in this study.

**Table S3.** Detection of variations in transcript isoform utilization by Majiq and Bambu-ISAR analysis. (A-C) Output of the MAJIQ algorithm (voila.tsv table) used to detect and quantify alternative splicing events on short read sequencing data, filtered by local splice variant (LSV) with a dPSI > 0.05 at least once. AS events were detected in LoVo cells infected by *Lm* for 2 h (A), 5 h (B) or 10 h (A) compared to non-infected cells and reported as delta in Percent Spliced-In (dPSI) per LSV. Data from 3 independent replicates at 0, 2, 5 hpi; 2 at 10 hpi. PSI, percent spliced-in. (D-F) Output of the Bambu-ISAR analysis. The R package Bambu was used to detect and quantify transcript variants in long-read data while R package ISAR was used to estimate the difference in usage of these isoforms across two conditions. Outputs give identifying information about the gene and isoform, the variation in both overall gene expression and isoform expression, the fraction of the total gene represented by each isoform in both conditions (isoform fractions, IF) and whether a premature termination codon (PTC) is present in the main expressed isoform. This analysis was used to compare isoform usage in LoVo cells infected by *Lm* for 2 h (D), 5 h (E) and 10 h (F) *versus* non-infected cells. Data from 3 independent replicates at 0, 2, 5 hpi; 2 at 10 hpi. FC, fold change; SE, standard error; PTC, premature termination codon. (G) Quantification of reads aligning with each one of the three possible local splicing variants at the 3’-junction of CIRBP exon 6, in non-infected cells and cells infected for 10 h by *Lm*. *Short-read analysis*. The number of reads aligning with the junctions of exon 6 with exons 7a or 7b was extracted from the outputs of MAJIQ; the number of reads spanning positions 1, 272, 051-1, 272, 052 was extracted from uniquely-mapped reads bam files for each sample. Each value was then normalized for library size. The proportion of each class with respect to the total number of normalized reads spanning this junction was then calculated for each condition. Fold-change (FC) values were calculated by dividing the number of normalized reads for each class in infected cells by their values in non-infected cells. *Long-read analysis*. Likewise, the number of long reads mapping to each one of the three types of junctions was extracted from Nanopore data and normalized for library size. The proportion of each class and FC were calculated from normalized counts as for short-read data. Data from 3 independent replicates at 0, 2, 5 hpi; 2 at 10 hpi. SD, standard deviation.

**Table S4.** CIRBP RIP-seq. (A-D) Output of DESeq2 analyses between the RNA fraction immunoprecipitated by CIRBP and either the input fractions in non-infected (NI) (A) and 10-h *Lm*-infected cells (B) or the IgG-immunoprecipitated fraction in NI (C) and *Lm* infected (D) cells. Result for each gene is reported as a log_2_ fold-change value and its corresponding adjusted *p*-value [*p*_adj_, DESeq false discovery rate (FDR)]. (E) For each CIRBP target, data of their enrichment in RIP-seq analysis and variations in expression assessed by RNA-seq, Ribo-seq and translational efficiency (TE) in 10-h infected (*Lm*) versus non-infected (NI) cells. (F) For each CIRBP target, data of their enrichment in RIP-seq analysis and variations in local splicing variants (LSVs) at the different junctions detected with MAJIQ for this target in 10-h infected (*Lm*) versus non-infected cells. FC, fold-change; SE, standard error; TE, translation efficiency; LSV, local splicing variation.

## Notes

https://www.ncbi.nlm.nih.gov/geo/query/acc.cgi?acc=GSE225417

https://www.ncbi.nlm.nih.gov/geo/query/acc.cgi?acc=GSE225516

https://www.ebi.ac.uk/ena/browser/view/PRJEB26593

### Competing Interest Statement

The authors have declared no competing interest.

### Summary of Updates

This version of the manuscript has been updated to incorporate novel experimental results regarding the CIRBP protein product, and strengthened correlative analysis of RIP-seq data with expression data.

## References

1. Fulda, S., Gorman, A.M., Hori, O. and Samali, A. (2010) Cellular stress responses: Cell survival and cell death. Int. J. Cell Biol., 2010, 214074.

2. Galluzzi, L., Yamazaki, T. and Kroemer, G. (2018) Linking cellular stress responses to systemic homeostasis. Nat. Rev. Mol. Cell Biol., 19, 731–745.

3. De Leeuw, F., Zhang, T., Wauquier, C., Huez, G., Kruys, V. and Gueydan, C. (2007) The cold-inducible RNA-binding protein migrates from the nucleus to cytoplasmic stress granules by a methylation-dependent mechanism and acts as a translational repressor. Exp. Cell Res., 313, 4130– 4144.

4. Zhong, P. and Huang, H. (2017) Recent progress in the research of cold-inducible RNA-binding protein. Future Sci. OA, 3, FSO246.

5. Corre, M. and Lebreton, A. (2023) Regulation of cold-inducible RNA-binding protein (CIRBP) in response to cellular stresses. Biochimie, 10.1016/j.biochi.2023.04.003.

6. Al-Fageeh, M.B. and Smales, C.M. (2009) Cold-inducible RNA binding protein (CIRP) expression is modulated by alternative mRNAs. RNA, 15, 1164–1176.

7. Haltenhof, T., Kotte, A., Bortoli, F.D., Schiefer, S., Meinke, S., Emmerichs, A.-K., Petermann, K.K., Timmermann, B., Imhof, P., Franz, A., et al. (2020) A conserved kinase-based body-temperature sensor globally controls alternative splicing and gene expression. Mol. Cell, 78, 57–69.e4.

8. Horii, Y., Shimaoka, H., Horii, K., Shiina, T. and Shimizu, Y. (2019) Mild hypothermia causes a shift in the alternative splicing of cold-inducible RNA-binding protein transcripts in Syrian hamsters. Am. J. Physiol.-Regul. Integr. Comp. Physiol., 317, R240–R247.

9. Neumann, A., Meinke, S., Goldammer, G., Strauch, M., Schubert, D., Timmermann, B., Heyd, F. and Preußner, M. (2020) Alternative splicing coupled mRNA decay shapes the temperature-dependent transcriptome. EMBO Rep., 21, e51369.

10. Lujan, D.A., Ochoa, J.L. and Hartley, R.S. (2018) Cold-inducible RNA binding protein in cancer and inflammation. WIREs RNA, 9, e1462.

11. Xia, Z., Zheng, X., Zheng, H., Liu, X., Yang, Z. and Wang, X. (2012) Cold-inducible RNA-binding protein (CIRP) regulates target mRNA stabilization in the mouse testis. FEBS Lett., 586, 3299–3308.

12. Gokhale, N.S., McIntyre, A.B.R., Mattocks, M.D., Holley, C.L., Lazear, H.M., Mason, C.E. and Horner, S.M. (2020) Altered m^6^A modification of specific cellular transcripts affects flaviviridae infection. Mol. Cell, 77, 542–555.e8.

13. Qiang, X., Yang, W.-L., Wu, R., Zhou, M., Jacob, A., Dong, W., Kuncewitch, M., Ji, Y., Yang, H., Wang, H., et al. (2013) Cold-inducible RNA-binding protein (CIRP) triggers inflammatory responses in hemorrhagic shock and sepsis. Nat. Med., 19, 1489–1495.

14. Ode, Y., Aziz, M., Jin, H., Arif, A., Nicastro, J.G. and Wang, P. (2019) Cold-inducible RNA-binding protein induces neutrophil extracellular traps in the lungs during sepsis. Sci. Rep., 9, 6252.

15. Celli, J. and Tsolis, R.M. (2015) Bacteria, the endoplasmic reticulum and the unfolded protein response: friends or foes? Nat. Rev. Microbiol., 13, 71–82.

16. Knowles, A., Campbell, S., Cross, N. and Stafford, P. (2021) Bacterial manipulation of the integrated stress response: A new perspective on infection. Front. Microbiol., 12.

17. Rodrigues, L.O.C.P., Graça, R.S.F. and Carneiro, L.A.M. (2018) Integrated stress responses to bacterial pathogenesis patterns. Front. Immunol., 9.

18. Kammoun, H., Kim, M., Hafner, L., Gaillard, J., Disson, O. and Lecuit, M. (2022) Listeriosis, a model infection to study host-pathogen interactions *in vivo*. Curr. Opin. Microbiol., 66, 11–20.

19. Pakos-Zebrucka, K., Koryga, I., Mnich, K., Ljujic, M., Samali, A. and Gorman, A.M. (2016) The integrated stress response. EMBO Rep., 17, 1374–1395.

20. Pillich, H., Chakraborty, T. and Mraheil, M.A. (2015) Cell-autonomous responses in *Listeria monocytogenes* infection. Future Microbiol., 10, 583–597.

21. Shrestha, N., Bahnan, W., Wiley, D.J., Barber, G., Fields, K.A. and Schesser, K. (2012) Eukaryotic initiation factor 2 (eIF2) signaling regulates proinflammatory cytokine expression and bacterial invasion. J. Biol. Chem., 287, 28738–28744.

22. Besic, V., Habibolahi, F., Noël, B., Rupp, S., Genovesio, A. and Lebreton, A. (2020) Coordination of transcriptional and translational regulations in human epithelial cells infected by *Listeria monocytogenes*. RNA Biol., 17, 1492–1507.

23. Gonzalez, M.R., Bischofberger, M., Frêche, B., Ho, S., Parton, R.G. and van der Goot, F.G. (2011) Pore-forming toxins induce multiple cellular responses promoting survival. Cell. Microbiol., 13, 1026–1043.

24. Pillich, H., Loose, M., Zimmer, K.-P. and Chakraborty, T. (2012) Activation of the unfolded protein response by *Listeria monocytogenes*. Cell. Microbiol., 14, 949–964.

25. Tattoli, I., Sorbara, M.T., Yang, C., Tooze, S.A., Philpott, D.J. and Girardin, S.E. (2013) *Listeria* phospholipases subvert host autophagic defenses by stalling pre-autophagosomal structures. EMBO J., 32, 3066–3078.

26. Vadia, S. and Seveau, S. (2014) Fluxes of Ca^2+^ and K^+^ are required for the listeriolysin O-dependent internalization pathway of *Listeria monocytogenes*. Infect. Immun., 82, 1084–1091.

27. Kwon, O.S., Mishra, R., Safieddine, A., Coleno, E., Alasseur, Q., Faucourt, M., Barbosa, I., Bertrand, E., Spassky, N. and Le Hir, H. (2021) Exon junction complex dependent mRNA localization is linked to centrosome organization during ciliogenesis. Nat. Commun., 12, 1351.

28. Vaquero-Garcia, J., Barrera, A., Gazzara, M.R., González-Vallinas, J., Lahens, N.F., Hogenesch, J.B., Lynch, K.W. and Barash, Y. (2016) A new view of transcriptome complexity and regulation through the lens of local splicing variations. eLife, 5, e11752.

29. Chen, Y., Sim, A., Wan, Y.K., Yeo, K., Lee, J.J.X., Ling, M.H., Love, M.I. and Göke, J. (2022) Context-aware transcript quantification from long read RNA-seq data with Bambu. 10.1101/2022.11.14.516358.

30. Vitting-Seerup, K. and Sandelin, A. (2019) IsoformSwitchAnalyzeR: analysis of changes in genome-wide patterns of alternative splicing and its functional consequences. Bioinformatics, 35, 4469–4471.

31. Eldridge, M.J.G., Cossart, P. and Hamon, M.A. (2020) Pathogenic biohacking: Induction, modulation and subversion of host transcriptional responses by *Listeria monocytogenes*. Toxins, 12, 294.

32. Lareau, L.F., Inada, M., Green, R.E., Wengrod, J.C. and Brenner, S.E. (2007) Unproductive splicing of SR genes associated with highly conserved and ultraconserved DNA elements. Nature, 446, 926–929.

33. Müller-McNicoll, M., Rossbach, O., Hui, J. and Medenbach, J. (2019) Auto-regulatory feedback by RNA-binding proteins. J. Mol. Cell Biol., 11, 930–939.

34. Risso, G., Pelisch, F., Quaglino, A., Pozzi, B. and Srebrow, A. (2012) Regulating the regulators: serine/arginine-rich proteins under scrutiny. IUBMB Life, 64, 809–816.

35. Yang, R., Zhan, M., Nalabothula, N.R., Yang, Q., Indig, F.E. and Carrier, F. (2010) Functional significance for a heterogenous ribonucleoprotein A18 signature RNA motif in the 3’-untranslated region of ataxia telangiectasia mutated and Rad3-related (ATR) transcript. J. Biol. Chem., 285, 8887– 8893.

36. Boehm, V., Kueckelmann, S., Gerbracht, J.V., Kallabis, S., Britto-Borges, T., Altmüller, J., Krüger, M., Dieterich, C. and Gehring, N.H. (2021) SMG5-SMG7 authorize nonsense-mediated mRNA decay by enabling SMG6 endonucleolytic activity. Nat. Commun., 12, 3965.

37. Li, J., Cai, Z., Vaites, L.P., Shen, N., Mitchell, D.C., Huttlin, E.L., Paulo, J.A., Harry, B.L. and Gygi, S.P. (2021) Proteome-wide mapping of short-lived proteins in human cells. Mol. Cell, 81, 4722–4735.e5.

38. Aziz, M., Brenner, M. and Wang, P. (2019) Extracellular CIRP (eCIRP) and inflammation. J. Leukoc. Biol., 106, 133–146.

39. Ribet, D., Hamon, M., Gouin, E., Nahori, M.-A., Impens, F., Neyret-Kahn, H., Gevaert, K., Vandekerckhove, J., Dejean, A. and Cossart, P. (2010) *Listeria monocytogenes* impairs SUMOylation for efficient infection. Nature, 464, 1192–1195.

40. Christie, M.P., Johnstone, B.A., Tweten, R.K., Parker, M.W. and Morton, C.J. (2018) Cholesterol-dependent cytolysins: from water-soluble state to membrane pore. Biophys. Rev., 10, 1337–1348.

41. Dal Peraro, M. and van der Goot, F.G. (2016) Pore-forming toxins: ancient, but never really out of fashion. Nat. Rev. Microbiol., 14, 77–92.

42. Krause, K.-H., Fivaz, M., Monod, A. and van der Goot, F.G. (1998) Aerolysin induces G-protein activation and Ca^2+^ release from intracellular stores in human granulocytes. J. Biol. Chem., 273, 18122–18129.

43. Hamon, M.A. and Cossart, P. (2011) K^+^ efflux Is required for histone H3 dephosphorylation by *Listeria monocytogenes* listeriolysin O and other pore-forming toxins. Infect. Immun., 79, 2839– 2846.

44. Ninomiya, K., Kataoka, N. and Hagiwara, M. (2011) Stress-responsive maturation of Clk1/4 pre-mRNAs promotes phosphorylation of SR splicing factor. J. Cell Biol., 195, 27–40.

45. de Oliveira Freitas Machado, C., Schafranek, M., Brüggemann, M., Hernández Cañás, M.C., Keller, M., Di Liddo, A., Brezski, A., Blümel, N., Arnold, B., Bremm, A., et al. (2023) Poison cassette exon splicing of SRSF6 regulates nuclear speckle dispersal and the response to hypoxia. Nucleic Acids Res., 51, 870–890.

46. Le Hir, H., Saulière, J. and Wang, Z. (2016) The exon junction complex as a node of post-transcriptional networks. Nat. Rev. Mol. Cell Biol., 17, 41–54.

47. Schroeder, A.L., Metzger, K.J., Miller, A. and Rhen, T. (2016) A novel candidate gene for temperature-dependent sex determination in the common snapping turtle. Genetics, 203, 557–571.

48. Kim, J.S., Jung, H.J., Lee, H.J., Kim, K.A., Goh, C.-H., Woo, Y., Oh, S.H., Han, Y.S. and Kang, H. (2008) Glycine-rich RNA-binding protein 7 affects abiotic stress responses by regulating stomata opening and closing in *Arabidopsis thaliana*. Plant J. Cell Mol. Biol., 55, 455–466.

49. Kim, J.Y., Kim, W.Y., Kwak, K.J., Oh, S.H., Han, Y.S. and Kang, H. (2010) Glycine-rich RNA-binding proteins are functionally conserved in *Arabidopsis thaliana* and *Oryza sativa* during cold adaptation process. J. Exp. Bot., 61, 2317–2325.

50. Nicaise, V., Joe, A., Jeong, B., Korneli, C., Boutrot, F., Westedt, I., Staiger, D., Alfano, J.R. and Zipfel, C. (2013) Pseudomonas HopU1 modulates plant immune receptor levels by blocking the interaction of their mRNAs with GRP7. EMBO J., 32, 701–712.

## References

1. Weinmaier, T., Riesing, M., Rattei, T., Bille, J., Arguedas-Villa, C., Stephan, R. and Tasara, T. (2013) Complete genome sequence of *Listeria monocytogenes* LL195, a serotype 4b strain from the 1983-1987 listeriosis epidemic in Switzerland. Genome Announc., 1, e00152–12.

2. Lecuit, M., Ohayon, H., Braun, L., Mengaud, J. and Cossart, P. (1997) Internalin of *Listeria monocytogenes* with an intact leucine-rich repeat region is sufficient to promote internalization. Infect. Immun., 65, 5309–5319.

3. Besic, V., Habibolahi, F., Noël, B., Rupp, S., Genovesio, A. and Lebreton, A. (2020) Coordination of transcriptional and translational regulations in human epithelial cells infected by *Listeria monocytogenes*. RNA Biol., 17, 1492–1507.

4. Glomski, I.J., Gedde, M.M., Tsang, A.W., Swanson, J.A. and Portnoy, D.A. (2002) The *Listeria monocytogenes* hemolysin has an acidic pH optimum to compartmentalize activity and prevent damage to infected host cells. J. Cell Biol., 156, 1029–1038.

5. Ribet, D., Hamon, M., Gouin, E., Nahori, M.-A., Impens, F., Neyret-Kahn, H., Gevaert, K., Vandekerckhove, J., Dejean, A. and Cossart, P. (2010) *Listeria monocytogenes* impairs SUMOylation for efficient infection. Nature, 464, 1192–1195.

6. Farrand, A.J., LaChapelle, S., Hotze, E.M., Johnson, A.E. and Tweten, R.K. (2010) Only two amino acids are essential for cytolytic toxin recognition of cholesterol at the membrane surface. Proc. Natl. Acad. Sci. U. S. A., 107, 4341–4346.

7. Iacovache, I., Paumard, P., Scheib, H., Lesieur, C., Sakai, N., Matile, S., Parker, M.W. and van der Goot, F.G. (2006) A rivet model for channel formation by aerolysin-like pore-forming toxins. EMBO J., 25, 457–466.

